# PanGIA: A Metagenomics Analytical Framework for Routine Biosurveillance and Clinical Pathogen Detection

**DOI:** 10.1101/2020.04.20.051813

**Authors:** Po-E Li, Joseph A. Russell, David Yarmosh, Alan G. Shteyman, Kyle Parker, Hillary Wood, J.R. Aspinwall, Richard Winegar, Karen Davenport, Chien-chi Lo, John Bagnoli, Phillip Davis, Jonathan L. Jacobs, Patrick S.G. Chain

## Abstract

Metagenomics is emerging as an important tool in biosurveillance, public health, and clinical applications. However, ease-of-use for execution and data analysis remains a barrier-of-entry to the adoption of metagenomics in applied health and forensics settings. In addition, these venues often have more stringent requirements for reporting, accuracy, and precision than the traditional ecological research role of the technology. Here, we present PanGIA (**<u>Pan</u>**<u>-</u>**<u>G</u>**enomics for **<u>I</u>**nfectious **<u>A</u>**gents), a novel bioinformatics analysis platform for hosting, processing, analyzing, and reporting shotgun metagenomics data of complex samples suspected of containing one or more pathogens. PanGIA was developed to address gaps that often preclude clinicians, medical technicians, forensics personnel, or other non-expert end-users from the routine application of metagenomics for pathogen identification. Though primarily designed to detect pathogenic microorganisms within clinical and environmental metagenomics data, PanGIA also serves as an analytical framework for microbial community profiling and comparative metagenomics. To provide statistical confidence in PanGIA’s taxonomic assignments, the system provides two independent estimations of probability for species and strain level detection. First, PanGIA integrates coverage data with ‘uniqueness’ information mapped across each reference genome for a stand-alone determination of confidence for each query sequence at each taxonomy level. Second, if a negative-control sample is provided, PanGIA compares this sample with a corresponding experimental unknown sample and determines a measure of confidence associated with ‘detection above background’. An integrated graphical user interface allows interactive interrogation and enables users to summarize multiple sample results by confidence score, normalized read abundance, reference genome linear coverage, depth-of-coverage, RPKM, and other metrics to detect specific organisms-of-interest. Comparison testing of the PanGIA algorithm against a number of recent k-mer, read-mapping, and marker-gene based taxonomy classifiers across various real-world datasets with spiked targets shows superior mean positive predictive value, sensitivity, and specificity. PanGIA can process a five million paired-end read dataset in under 1 hour on commodity computational hardware. The source code and documentation are publicly available at https://github.com/LANL-Bioinformatics/PanGIA or https://github.com/mriglobal/PanGIA. The database for PanGIA can be downloaded from ftp://bioinformatics.mriglobal.org/. The full GUI-based PanGIA analysis environment is available in a Docker container and can be installed from https://hub.docker.com/r/poeli/pangia/.

## INTRODUCTION

Unbiased whole-metagenome shotgun (WMS) sequencing has emerged as a promising solution for rapid pathogen identification in various applied settings, from the clinic[1] to public health, food-safety[2], and biodefense[3], due to rapidly declining costs and increasing accessibility of sequencing technologies in recent years. A primary challenge in applying this technology is taxonomic profiling – a process to assign taxonomy classification accurately – which requires high species-level accuracy and a statistically sound method to interpret the assignment results. In non-research settings, such as clinical and biosurveillance applications, there are numerous additional challenges such as ‘ease-of-use’ and analytical speed on commodity computers[4].

To address the taxonomic profiling challenge, an array of software tools and algorithms have been developed in the past[5-7]. Those algorithms can be broadly categorized into two groups of tools, each of which require the pre-construction of a tool-specific database before a search is conducted. The read-mapping approach aligns sequencing reads directly to a reference database, while the k-mer counting approach counts the k-mers in reads and compares them to the k-mer counts collected from a database of reference genomes. Examples of alignment methods include BLAST[8], BWA[9], BowTie2[10], and DIAMOND[11]. While the alignment-based approach has shown high sensitivity in metagenomic profiling[5-7], GOTTCHA[12], MetaPhlAn2[13] and mOTUs[14] improved specificity by leveraging hierarchical genomic marker-based databases. Despite its well-balanced specificity and sensitivity, alignment-based approaches against large databases are relatively slow since the process requires aligning all reads to the entire reference genome database. Other recent tools have devised methods to improve speed by constructing a database of annotated protein-coding genes from microbial genomes (Kaiju[15]) or implementing a streamlined database that eliminates identical sequences between strains within a species (Centrifuge[16]). Kaiju and Centrifuge implement the Ferragina-Manzini (FM) Index, a technique that constructs a ‘look-up’ table for occurrence counts of each characteristic (e.g., protein-coding genes or unique genomic loci). K-mer-based approaches were developed to avoid the alignment process. This technique matches k-mers from reads to that of a database of k-mers derived from reference genomes with more than a tenfold speed gain compared with alignment-based approaches. Some example k-mer counting tools include Kraken[17], LMAT[18], and CLARK[19]. Selecting the right size of *k* for building k-mers remains a tradeoff between sensitivity and specificity and defaults to different values in different software, (e.g. *k=*31 in Kraken and CLARK while LMAT uses *k=*20). Running k-mer counting tools requires loading the entire k-mer database into computer memory, which often takes over 70 GB of RAM, thus limiting its application in many settings outside of a research institution. MetaOthello[20] lowers the memory requirement to less than 32 GB by utilizing an optimized data structure. A new addition to these two approaches is a pseudo-alignment approach, which can rapidly determine the compatibility of reads with a reference database indexed using a de Bruijn Graph. Some example tools include Kallisto[21] and Karp[22]. k-SLAM[23] utilizes both local pseudo-alignment and pseudo-assembly to improve the taxonomic profiling of multi-mapping reads that can be aligned to different reference genomes. More details about different approaches in taxonomic profiling can be found in recent reviews [24].

Associated with some of the software tools, various methods have been developed to provide confidence metrics for sequence read alignments and taxonomy classifications. For example, Kraken and CLARK apply a confidence score for each read using the ratio of unique to ambiguous k-mers. Other methods, GAAS[25] and GRAMMy[26] implement a confidence metric by using iteratively evolving abundance estimations with an average genome length, Expectation-Maximization (EM) algorithm, and genome similarity implied by read-alignment. GASiC[27] furthered these efforts by incorporating a pre-calculated whole genome similarity matrix of all references and performing a statistical test with a bootstrapping procedure. However, an even-weighted pairwise similarity matrix could reduce the strength of signals generated by reads aligned to unique regions of a genome, which would be problematic for the differentiation of closely-related genomes. Thus, while confidence scoring remains a desirable feature of taxonomy profilers, no clear method has emerged.

An ideal computational tool for applied metagenomic profiling in clinical and/or biosurveillance settings needs to address the following: 1) a robust balance of sensitivity and specificity, 2) a comprehensive database of full, high-quality genomes, 3) a confidence metric that can resolve, or at least inform, strain-level detection, 4) a confidence metric that can resolve detected pathogens from contaminating DNA and/or background microbial communities, 5) a robust workflow with a graphical-user-interface (GUI) for non-experts that presents results in a clear, concise manner and enables rapid discernment of organisms-of-interest, 6) the ability to run on commodity computational hardware (i.e., < 32GB RAM), and 7) remain agnostic of sequencing platform. None of the metagenomics taxonomy classifiers discussed above, or published to date, meet all of these objectives.

Here, we present PanGIA (Pan-Genomics for Infectious Agents), a novel read-alignment based metagenomics taxonomy classification tool. PanGIA delivers comprehensive metagenomic data analysis (organism identification with confidence scoring of taxonomic hits, together with other metrics), coupled with an interactive user-interface that allows dynamic exploration of the evidence underlying identified pathogens (and other organisms). PanGIA addresses some of the above-discussed barriers to the adoption of sequencing in applied settings. We utilize taxonomy-aware unique genome sequence signatures, developed as part of GOTTCHA[12], as a basis for the development of a novel confidence scoring system to report the presence of identified organisms. We validated PanGIA profiling results using a series of *in silico*-derived datasets, and benchmarked it against a number of recently published or updated tools. We further tested and evaluated its use on three different sets of clinical samples, as well as previously published clinical data from validation of other taxonomy tools, to demonstrate its real-world utility. The source code and documentation are publicly available at https://github.com/LANL-Bioinformatics/PanGIA or https://github.com/mriglobal/PanGIA. The database for PanGIA can be downloaded from ftp://bioinformatics.mriglobal.org/. The full GUI-based PanGIA analysis environment is available in a Docker container and can be installed from https://hub.docker.com/r/poeli/pangia/.

## MATERIALS AND METHODS

### Reference database and custom taxonomy construction

The PanGIA database includes three major components: genome sequences, their taxonomic lineage information, and pre-calculated coordinates of genome uniqueness at each taxonomic level. The genome sequences database includes bacteria, archaea, and virus genome entries that are categorized as representative and/or reference genomes in NCBI RefSeq (release 89)[28]. In addition, it also includes all complete genomes of CDC bioterrorism agents (<u>Category A, B and C</u>)[29] and *Plasmodium* species (causative agent of malaria). Common host and vector genomes (e.g. human and mosquito genomes) are also included to allow for *in-silico* de-hosting, prior to read classification. Additional host genomes can be added for customized de-hosting, for example in veterinary medicine applications. In the case where more than one isolate exists for a given strain, only one representative genome sequence is used. Additional information, including genome length, taxonomy ID and pre-defined tags were added to the FASTA header of the sequences before they were indexed using BWA.

PanGIA relies on the NCBI Taxonomy Database[29] to provide taxonomic lineage information. As needed, custom taxonomic identifiers were assigned to strains by appending unique suffixes to its closest taxonomy node. For instance, the first strain of *Vibrio cholerae* (taxonomy ID 666) with no strain-level taxonomy ID is assigned to 666.1

PanGIA uses taxonomy-aware genome uniqueness in order to calculate a confidence metric that is independent from read abundance. A genome uniqueness table was generated using the database construction algorithm accompanying the GOTTCHA taxonomy profiler, in order to identify unique regions within all genomes at all eight levels of taxonomy[12].

### Datasets used for Benchmarking and Confidence Scoring

For benchmarking of taxonomic tools, two separate types of *in-silico* datasets were generated. One type was to determine traditional metrics of analytical sensitivity and specificity, and another type was to measure ‘near-neighbor’ specificity that is unique to the metagenomic taxonomy classification problem. Traditional analytical sensitivity and specificity are measured from the presence/absence of ‘targets’ in known target-positives vs. known target-negatives, to produce values for *true positives* (TP), *true negatives* (TN), *false positives* (FP), and *false negatives* (FN). Analytical sensitivity is then calculated as (TP)/(TP+FN). Analytical specificity is calculated as (TN)/(TN+FP). When using these metrics for benchmarking of metagenomics taxonomy classifiers, the level at which off-target near-neighbor species are detected from a given taxonomy classifier is not captured. For example, if *Staphylococcus aureus* is in a clinical sample and a sequence read-based signal from that organism is acquired in the clinical sample, then that is a *true positive*. If mis-classified *Staphylococcus epidermidis* reads (that actually originated from the etiological *S. aureus*) are also indicated for the clinical sample, this indication is **not** captured as a *false positive* by the traditional metrics listed above. Thus, we constructed two datasets; one that would allow us to measure traditional sensitivity and specificity, and one to measure off-target near-neighbor mis-classification.

For the measurement of analytical sensitivity and specificity across all tools, we selected 52 organisms (24 bacteria and 28 viruses, **Supplementary File 1**) representing a range of Gram-positive and Gram-negative bacterial pathogens, as well as RNA and DNA viral pathogens. Starting from reference genomes of each pathogen, we used a custom script, combined with DWGSIM[30] (**Supplementary File 1**), to generate triplicate sets of 100, 1,000, 10,000 and 100,000 2×75 bp paired-end *in-silico*-simulated Illumina reads per organism. Reads at each depth-level were combined and ‘read-replaced’ within a 10 million read dataset from actual experimental sequencing of a dirty forensic swab rubbed through 5 g of reference background material (loam soil, Sigma-Aldrich, St. Louis, MO) that comprised ∼30M reads in total. For example, at the lowest ‘titer level’, there were 5,200 reads (100 reads per organism x 52 organisms) *in-silico* target reads. 5,200 random reads were subtracted from the 10M read dataset and replaced with the 5,200 target reads for a total of 10 million paired-end reads with each target representing 0.001% of the total. Individual target abundance increased to 0.01%, 0.1%, and 1% in the higher ‘titer level’ datasets. Each titer level was independently produced three times and analyzed independently with each classifier for triplicate data points for each target at each titer level. Triplicate negative controls of ‘unspiked’ reference background material datasets (10 million reads each, randomly sub-sampled from the total 30M read set) were also processed with each classifier.

For the measurement of off-target near-neighbor mis-classifications, we created ‘naked read datasets’ for four pathogens – *Burkholderia mallei, Staphylococcus aureus*, Venezuelan Equine Encephalitis virus, and Variola virus (Gram neg. bacteria, Gram pos. bacteria, RNA virus, DNA virus, respectively). ‘Naked read datasets’ are *in silico-*generated reads that are not combined with any real-world sequencing data, and were run by themselves through each taxonomic classifier to assess incorrect classifications. An ‘ideal’ taxonomy classifier, when given 1,000 reads of *S. aureus* (for example), should return 1,000 reads of <u>only</u> *S. aureus*, and 0 reads for any other organism. Of course, due to evolutionary sequence homology, such a perfect classifier is not possible – there is simply too much similarity between conserved sequences in related clades, leading to reads being ‘multi-mapped’, or assigned to multiple taxa despite originating from only one actual organism in a sample. Especially in applied settings, the degree to which ‘multi-mapping’ manifests itself and muddles the interpretation of single-target presence in community-level data is an important quality to determine. However, it is difficult to quantify from community-level data. Thus, to more effectively query this important metric, we created the ‘naked read’ datasets in triplicate for 5 signal intensities – 10, 100, 1,000, 10,000, and 100,000 reads. The number of reads recovered at the species level for each organism and the number of off-target near-neighbor mis-classifications at the species level were recorded.

To test both standalone and background confidence scoring behavior, we generated a ‘real-world’ sequencing dataset across various titers of the microbiome standard ATCC^®^ MSA-2002™ 20-Strain Even Mix Whole Cell material that were spiked into forensics swab matrices that had been rubbed through reference background soil material (Sigma-Aldrich, MO, USA). The ATCC material was diluted to 1e3, 1e5, and 1e6 CFU/ml, spiked into the soiled swab matrices, and extracted, processed, library-prepared, and sequenced with the Sample-2-Sequence (S2S) metagenomics analysis workflow (see *Methods*, Parker et al. 2019). To determine real-world vs. idealized performance, we analyzed each of the S2S-processed microbiome standard samples with PanGIA. For each of the 20 organisms that were expected, we determined the number of reads mapping at the species-level and, using the sequence read IDs from the SAM file, we removed the corresponding reads from the data and replaced them with a matching number of DWGSIM-generated *in silico* reads for each organism. The *in silico* reads are an ‘ideal’ sampling of the genome and not subject to any biases that may exist in ‘real world’ data (e.g., high copy-number rRNA, etc.). Both standalone score and background score were recorded for each of the 20 organisms at each titer level across the ‘real-world’ and *in silico* data.

Finally, as a real-world example, human clinical samples were tested. Two were extracted from nasopharyngeal swabs and one from serum. The nasopharyngeal swabs were collected from one patient infected with influenza virus one patient infected with *Bordetella pertussis*. The serum sample was from a patient infected with Dengue virus. All samples and clinical lab analyses were provided by LANL Occupational Medicine and New Mexico Department of Health. All samples were sequenced on Illumina’s NextSeq platform with 2×151 bp v2 chemistry and generated 10,655,304 reads for the influenza sample, 85,775,914 reads for the *B. pertussis* sample, and 7,295,652 reads for the Dengue virus sample.

For all DWGSIM-generated *in-silico* reads used in this study, an Illumina error model (0.001-0.01) was used. All *in-silico* datasets used for comparative analyses are publicly available in the Dryad data repository, at the following DOI; 10.5061/dryad.hdr7sqvf1. The human clinical datasets are available in the NCBI’s Sequence Read Archive (SRA) under NCBI BioProject PRJNA594744.

### PanGIA taxonomy classification tool

To identify the organisms present within a sequencing dataset, PanGIA utilizes the BWA-MEM alignment algorithm to align reads to a reference genome database. The default aligner is BWA-MEM, although any read aligner that supports tracking read placements can be used. The GOTTCHA database construction algorithm was used to identify unique sequence coordinates for each reference genome at all eight major taxonomic ranks, from strain to superkingdom. Alignment results are combined with taxonomy/genome uniqueness of the hits to both filter the results and provide PanGIA’s ‘standalone’ (SA) confidence score at the appropriate taxonomic level. In addition, PanGIA calculates a ‘background’ (BG) confidence score associated with the statistical confidence that a detected taxa is present at significant levels above those observed in a pre-PanGIA-analyzed laboratory negative control, or ‘background’ sample. The specifics of standalone and background confidence score calculation can be found in the *Results* section.

PanGIA aligns short reads to the BWA-indexed sequence database with “-k40 -T60 -H150 -B2” options, and reports multiple hits. For the *i*^*th*^ mapped read, *r*_*i*_, up to 30 qualified hits (*h*_*i,j*_) that align to different genomes (*j*) and have an equivalent maximum alignment score will undergo downstream processing, such that 1 ≤ |*h*_*i*_| ≤ 30. A taxonomic specificity level *s*_*i*_ of *r*_*i*_ is defined using the taxonomic rank of the last/lowest common ancestor (LCA) of all qualified hits, where *s*_*i*_ ∈ {strain, species, genus, …, superkingdom}.

PanGIA tracks both raw and normalized read counts on a per organism basis for all eight taxonomic levels. For every organism *j*, the raw read count (*R*) is the total number of reads that hit to any of its replicons (contigs, chromosomes and plasmids), such that 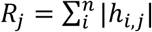. We normalize *R* based on the number of organisms the read hits (NR; i.e., dividing by the number of hits), the mapping percent identity (NI) and the combination of both (NB). For each organism *j*, the total hits normalized by *NR* are: 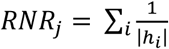, *where* 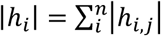 and |*h*_*i,j*_| · *c*_*i*_. The NI normalized organism read-count multiplies (normalizes by) the percent identity for each read: 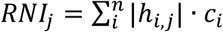, where *c*_*i*_ is the mapping identity. Combing both NR and NI, the NB normalized read count for organism *j* is 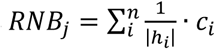.

Other than counting reads, PanGIA also tracks the average depth of coverage (*DC*) of the organism *j*: 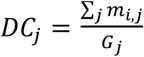, where *m* is the total mapped bases of the i^th^ read that hits genome *j* and *G*_j_ is the genome size of the genome *j*. NR normalization is also applied to *DC* as 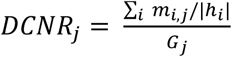. The raw and normalized *Rs* and *DC*s are further summed up to other major taxonomies all the way to superkingdom.

PanGIA also provides a taxonomy rank-specific (RS) version of *R*s and *DC*s that only takes into account reads that specifically map to genomes within a given taxonomic level. For instance, the *RS-DCNR* for taxon *t* is the DCNR calculated by reads that hit genomes belonging to taxon *t* only. The SAMtools[31] depth command is used to calculate the depth information by leveraging all mapped reads classified as more specific than class-level (*s*_*i*_ ∈ {strain, species, genus, …, order}). The linear length (L_k_) is defined as the number of mapped bases in the organism *j*. The genome coverage is: 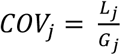. PanGIA reports the filtered profiling results using adjustable defaults of: minimum 10 reads, minimum 2.5 rank-specific normalized reads, minimum 0.4% genome coverage, minimum 1% depth of coverage and minimum 0.09% rank specific depth of coverage normalized references.

### Comparative metrics

Youden’s J[32] and F1 scores were used to evaluate and compare the performance of PanGIA with other tools. True positives (TP) were defined as read-based detection of truly present target organisms at the species level in each spiked dataset (*assuming detection above each tool’s implemented cut-off criteria, but otherwise regardless of read-count*). False negatives (FN) were defined as *missing* read-based detection and taxonomic-classification of truly present target organisms at the species level in each spiked dataset. True negatives (TN) were defined as **no** read-based detection and taxonomic classification of target organisms at the species level in ‘unspiked’ datasets. False positives (FP) were defined as read-based detection and taxonomic-classification of target organisms at the species level in each ‘unspiked’ dataset, regardless of read-count. Analytical sensitivity was calculated as (TP)/(TP+FN). Analytical specificity was calculated as (TN)/(TN+FP). Youden’s J can be calculated: Y_j_ = (sensitivity + specificity) – 1. F1 score is measured as F_1_ = ((sensitivity^-1^ + precision^-1^)/2)^-1^, where precision = TP/(TP + FP). PanGIA was benchmarked against Centrifuge, CLARK-S[33], GOTTCHA, Kaiju, Kraken2, KrakenUniq[34], and Metaphlan2. Job runtime parameters for each tool are listed in **Supplementary File 1**.

### Interactive use of PanGIA and data visualization

We developed an interactive web-based visualizer, Interactive Metagenomic Taxonomy Viewer (IMTV), to display results using Bokeh[35]. Bokeh is a high-performance interactive visualization library to create plots, dashboards, and applications for presenting large datasets. PanGIA launches a custom Bokeh server process that routes data to the BokehJS client library at the front-end, dynamically using Tornado’s web framework.

The tab-delimited result file is used to visualize the integrated taxonomic profiles within a dashboard and a dot plot. Separately, individual genomes and their calculated coverage are displayed. The samtools-depth files are used to display the genome coverage plot at the strain level. The per-replicon basis depth files generated by the PanGIA pipeline are merged into single files for each organism. To create meaningful representations for large genomes, and avoid over-representation, the merged depth files are evenly resampled at up to 1500 positions. To allow the display of annotations (e.g. display of special marker genes) within the genome, a feature table with genome coordinates can be used to annotate the genome plot.

For the available genomes, we collected a pathogen list composed of pathogen names, taxonomies and some metadata that includes known hosts, sources of the sample, disease names, etc. These metadata are compiled from authorities, such as the United States CDC and the WHO. This table is used within the web GUI to display this information in a table format under the genome plot.

### PanGIA’s Select Agent Pathogen Markers

Species-specific markers were generated using T-MArC (https://github.com/mriglobal/T-MArC) for all biological select agents. T-MArC leverages the ShortBRED[36] algorithm to identify portions of genes that are unique. Markers are drawn from publicly available references and protein sequences on RefSeq and take the form of amino acid strings. Markers produced by ShortBRED are compared to all available references of the target agent as well as its near neighbors. Those markers that are found to be completely *exclusive* of closely related off-target species, and completely *inclusive* of the species-of-interest, are then ranked by how comprehensively they cover all strains of the target agent. These markers are binned according to this level of completeness, and these are included as available features to examine in the genome coverage plot.

### Code Availability

The source code and documentation are publicly available at https://github.com/LANL-Bioinformatics/PanGIA or https://github.com/mriglobal/PanGIA. The database for PanGIA can be downloaded from ftp://bioinformatics.mriglobal.org/. The full GUI-based PanGIA analysis environment is available in a Docker container and can be installed from https://hub.docker.com/r/poeli/pangia/.

## RESULTS

### PanGIA taxonomy profiler overview

The PanGIA pipeline begins with aligning input reads to all provided bacterial, viral and host genome sequences (**Figure 1**). Instead of simply identifying all genomes that sequence reads are mapped to, we focus on how specifically reads are aligned. Thus, PanGIA provides not only a simple read count and depth of coverage for each organism, but also reports how many mapped reads hit equally well to other organisms, the percent identity of the hits, and from these, derives a ‘standalone’ (SA) confidence score (i.e., a score that does not need control, reference samples). If a control sample is available, PanGIA can also calculate the background community profile (by calling up pre-analyzed PanGIA results on a negative control in JSON format) to provide a separate, ‘signal-above-background’ confidence score. Due to the sequencing platform and the variety of host genomes (human or others) and the diversity therein, it is difficult to find an optimized suite of parameter settings for removing host reads using a mapping method. Thus, all reads are mapped to both target and host genomes. Reads that map better to the host versus any target reference genome are discarded.

**Figure 1:**
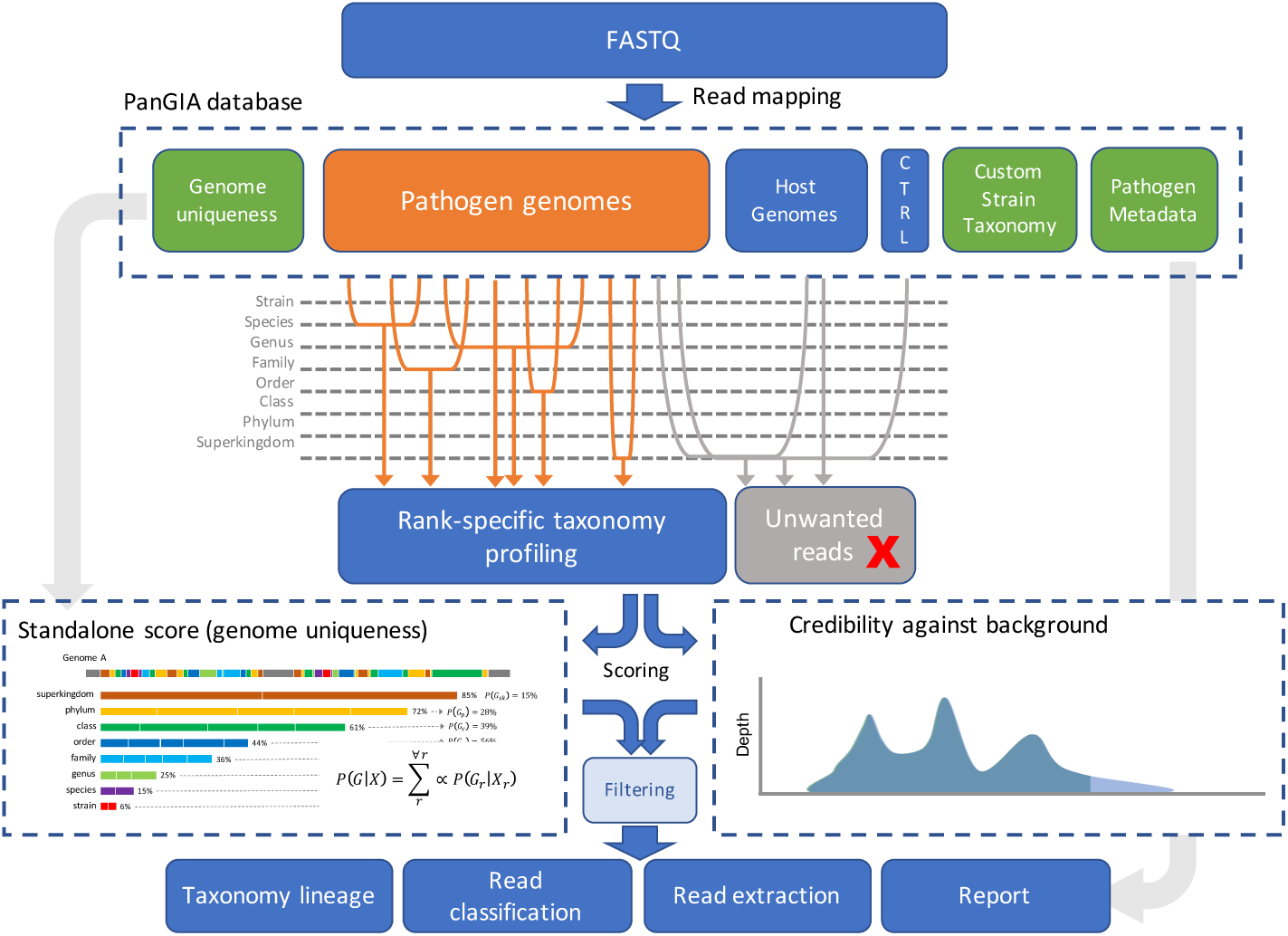
Workflow of PanGIA taxonomy profiler.

### Developing a novel standalone confidence metric

To provide a confidence metric of whether mapped genomes are in fact found within a sample, we propose a hierarchical Bayesian model to evaluate the credibility of read matches by gathering evidence from the mapping specificity of each read at each taxonomy level (**Figure 2**).

**Figure 2:**
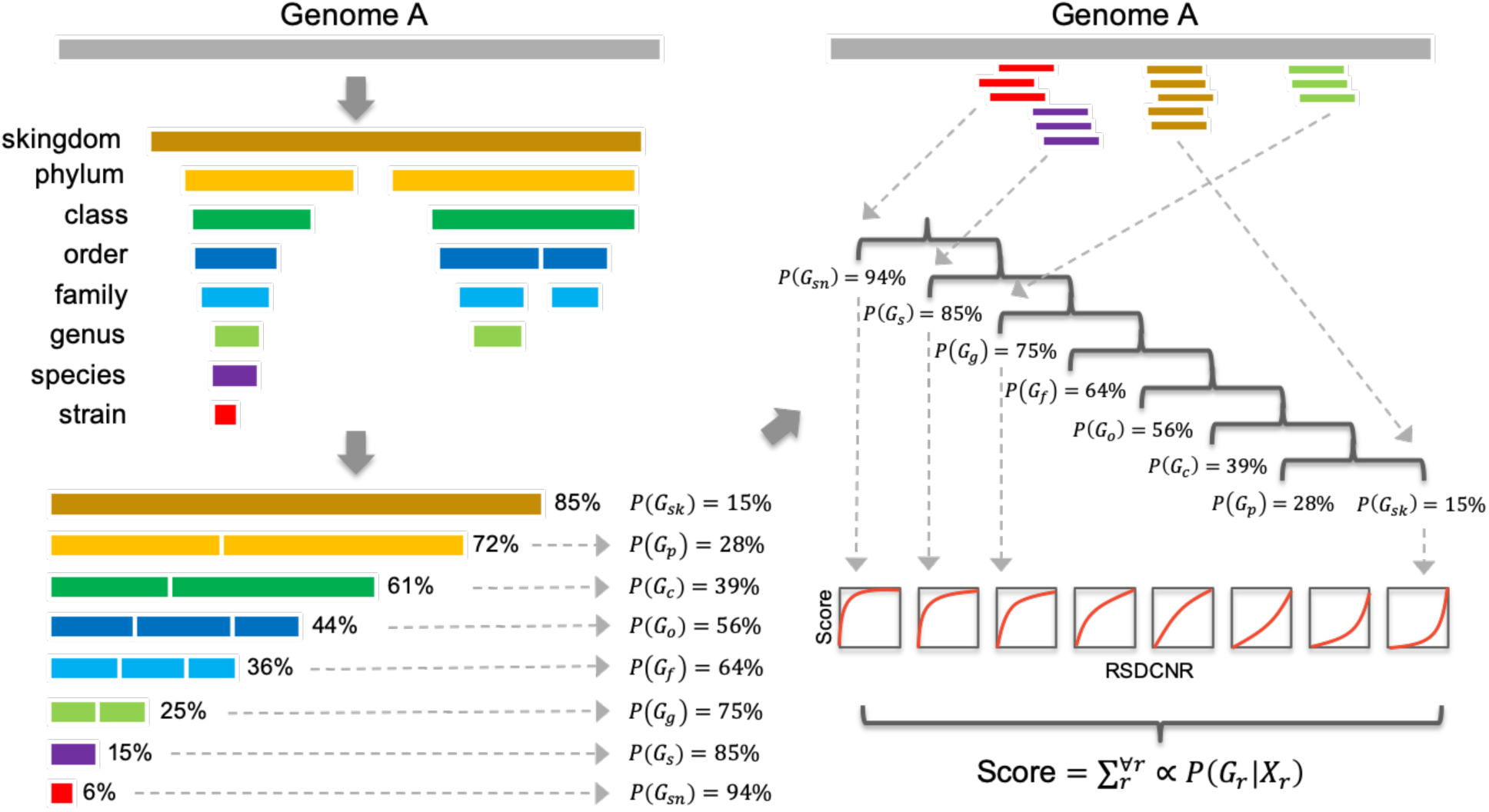
Overview of standalone scoring. The standalone score is calculated by cross-referencing the depth of coverage across a reference genome with the amount of that genome that is informative to a particular taxonomic level. Each reference genome in PanGIA’s database has the ‘uniqueness’ modeled against the rest of the genomes in the database. This coverage information and ‘uniqueness’ information is then translated to a probability of correct assignment at the particular taxonomic level.

Given the GOTTCHA database of unique genomic regions, and the full taxonomy of database entries, we are able to ascertain what fractions of the unique genome are covered at each taxonomic level and use a posterior probability of what would be expected if the genome was randomly sampled. Let *X* denote the number of reads that map to taxon *G* (specific organism’s genome) and *X*_*r*_ is the number of reads that map to *G* with specificity-level at *r* rank. The probability of *G* existing in the sample can be inferred to the posterior probability *P*(*G*_*r*_|*X*_*r*_), where r belongs to one of the eight major ranks from strain to superkingdom. Based on Bayes’ theorem, this can be expanded to:

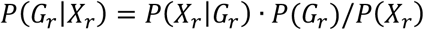

Even though reads can map uniquely to taxonomy *G*, there remains some uncertainty due in part to the incompleteness of our reference databases – reads could map equally well to a genome that may not be included in our database and has yet to be sequenced. By utilizing only the unique region(s) in each genome for each level of taxonomy, we remove the ambiguity of many reads that can be mapped equally well to multiple genomes. We assign the “certain” proportion of the genome, denoted as *p*_*r*,_ according to the uniqueness matrix to the priors *P*(*G*_*r*_). The normalized rank-specific depth-of-coverage (RS-DCNR) is assigned to the probability of observing *X*_*r*_ given the taxonomy is present:

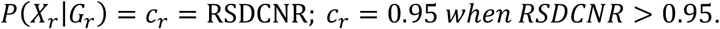

The *P(X*_*r*_ ) can be rewritten and simplified:

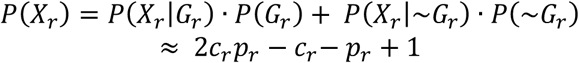

Summing up the weighted values from each rank, a **standalone score** is represented by the following formula, given *r* belongs to a specific rank, and up to superkingdom:

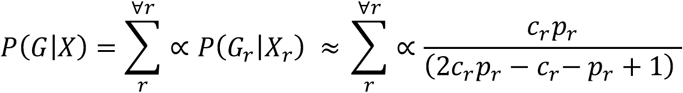

### Developing a background, control-based confidence metric

An alternative confidence metric that can be applied for the purpose of identifying outliers, is to compare the results of a target sample with one or more background (or control) samples, if available. Control samples are meant to represent what would typically be found in a sample, and can serve as what regularly occurs in the background (e.g., serum from a healthy donor when surveying for etiological agents from a patient with a fever of unknown origin, or better, a sample from the same patient prior to infection). Alternatively, a ‘laboratory blank’ can be processed to mask out common laboratory contaminants. The control sample(s) are intended to be run through PanGIA prior to testing any target samples. These are similarly analyzed against the available reference database to identify the background signal. By loading the output file of SAMtools Depth, the base-by-base positional information is flattened into binary masks that indicate whether sequences are present in the background community. These masks are saved to JSON files (mask files) and can be retrieved when running a target dataset. PanGIA will calculate a positional array that removes the depth information from the positions according to the mask file(s). For each genome of interest, background confidence scores are defined as the ratio of the sum of pre-masked depths to the sum of masked depths. A higher background confidence score for any given organism indicates a smaller portion of that genome was identified in the control sample that overlapped with that identified in the target sample (**Figure 3**).

**Figure 3:**
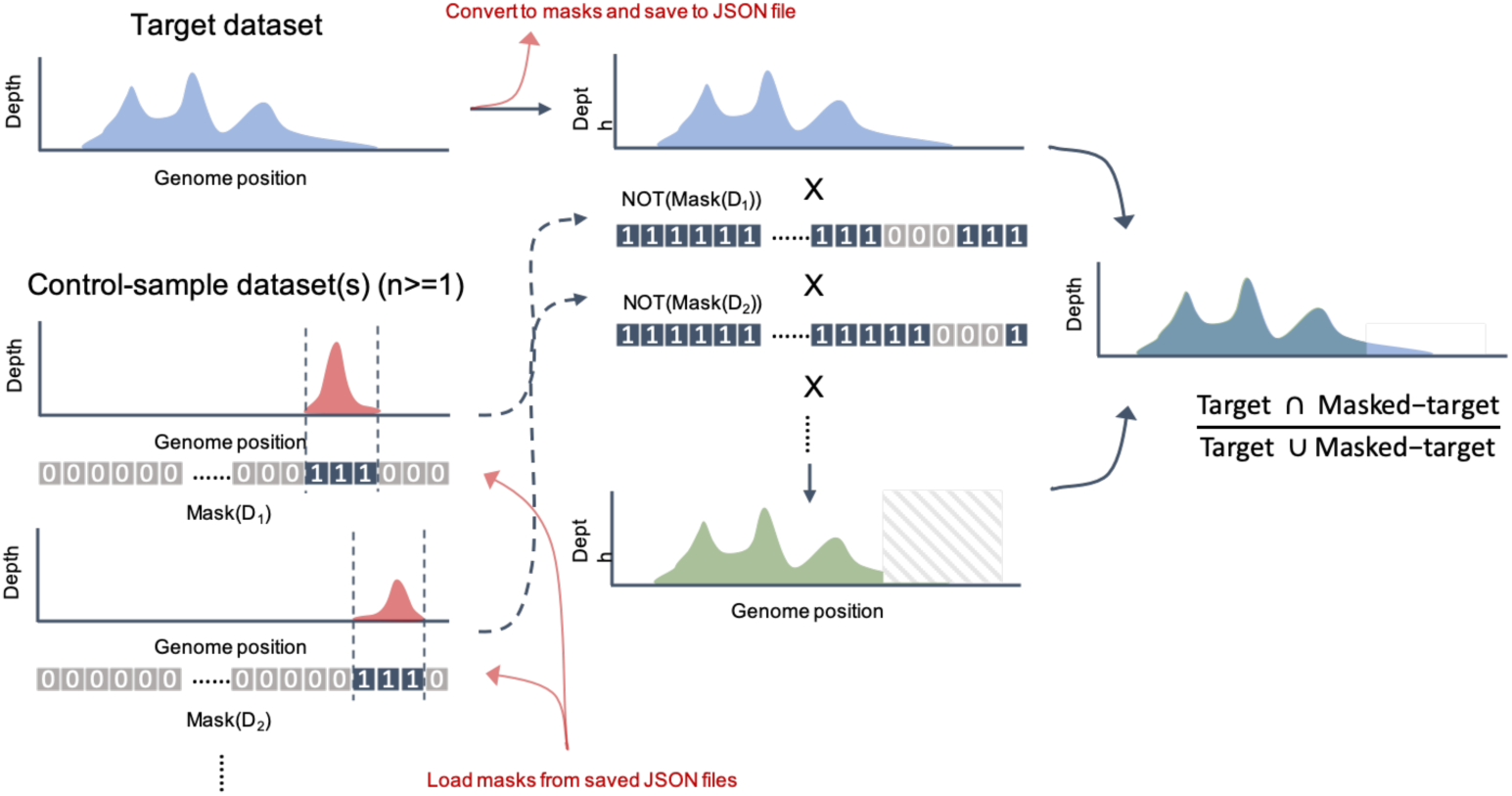
Overview of background confidence scoring. Genome coverage for target organism in analytical sample (blue) is overlaid with genome coverage in background control sample (red). Regions that overlap are ‘masked’ (hashed box) and the background confidence score is calculated by the intersection over the union of the two distributions (green over blue).

Incorporating two confidence metrics, that look at the data from separate but complementary angles, is a novel strategy that can aid in the interpretation of an individual organism’s presence, despite operational biases in real-world sequencing workflows. This can be observed by examining the distribution of both standalone and background confidence scores across *in-silico* and ‘real-world’ sequencing datasets. *In-silico* data, very often used in metagenomics taxonomy benchmarking studies, represents an ‘ideal’ sampling of each individual reference genome from all the organisms comprising a synthetic community dataset. The reference genomes are evenly sampled across their length at random, and the biases of actual operational workflows are not captured. This phenomenon is seen in **Figure 4**. To evaluate these issues, a mock community (ATCC^®^ MSA-2002™ 20-Strain Even Mix Whole Cell material) was extracted, processed and sequenced on an Illumina MiSeq machine, across a range of titer levels. The resulting data was analyzed with PanGIA. The number of reads associated with each organism in the real-world data were re-capitulated *in-silico* and also analyzed with PanGIA. In the ideal-sampling of the *in-silico* analysis, a progressively tighter clustering of high standalone and background confidence score is seen as the titer level increases, whereas a larger spread of both standalone and background scores is observed in the ‘real-world’ data. Some organisms with particularly crowded clades (e.g., *Clostridium Rhodobacter*) are difficult to classify with high-confidence at the species-level, even with *in-silico* data. However, the distinction between ‘ideal’ sampling and real-world sampling is evident for other organisms (e.g., *Bacillus spp., Bacteroides spp.*, etc). Most organisms converge towards high scores in both categories as titer level increases (at varying rates). Viral targets are observed to converge more quickly due to lower genome complexity, size, and lower chance of background presence, whereas bacterial targets with larger and more similar genomes, and higher likelihood of near-neighbor representation in background, converge more slowly and are more likely to have outliers.

**Figure 4:**
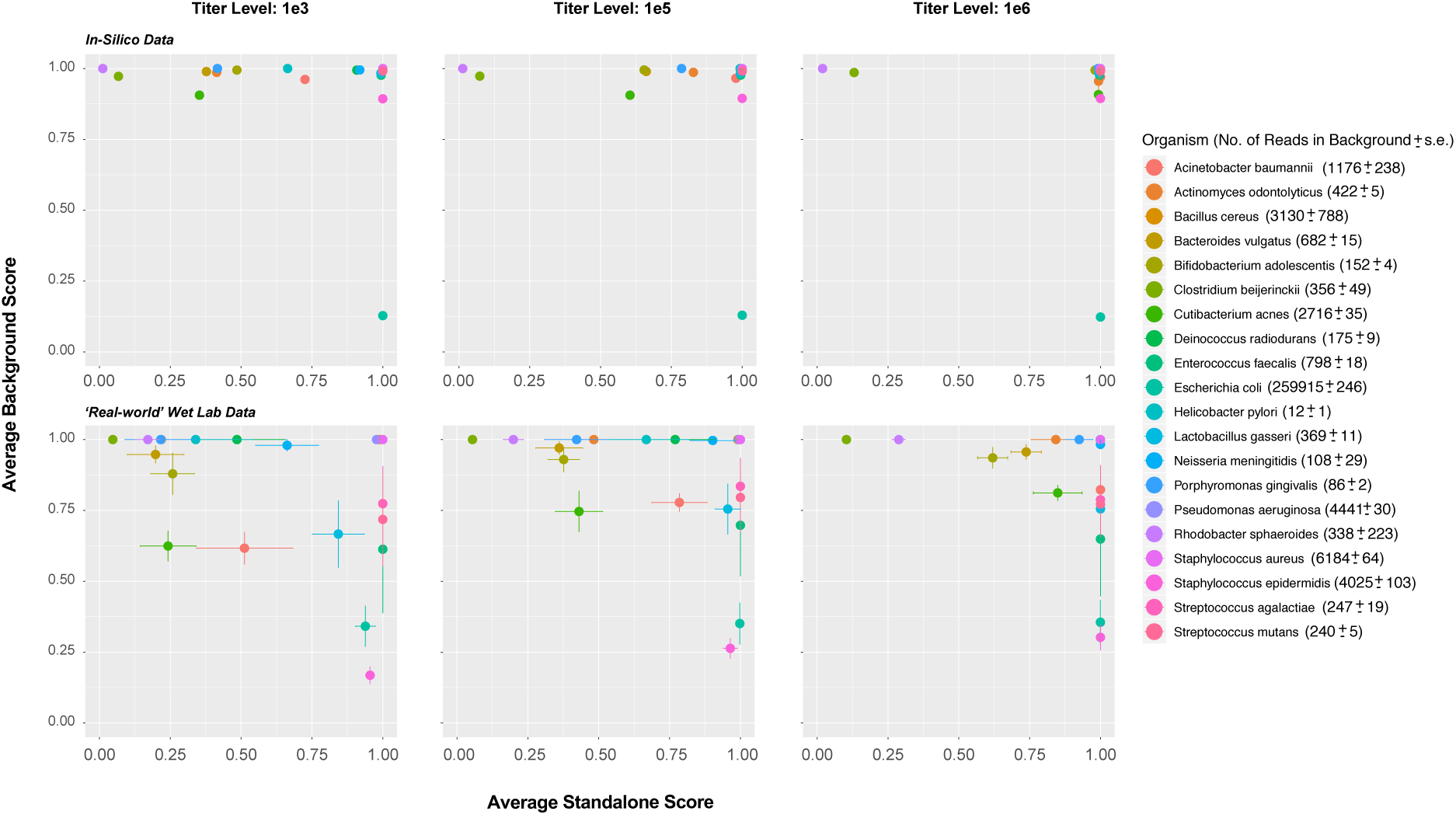
Relationship between standalone and background-control scoring across a range of mock titer levels in analysis of a mock community (ATCC® MSA-2002™ 20-Strain Even Mix Whole Cell material) from real wet-lab processing of the community (lower panels) and an in-silico re-capitulation of the community data (top panels).

### Comparison among different classifiers

We analyzed PanGIA against a range of other tools (Centrifuge, CLARK-S, GOTTCHA, Kaiju, Kraken2, KrakenUniq, Metaphlan2) under a community-profiling context, with precisely known read-level quantities of 52 target bacteria and viruses amongst a reference soil-matrix background community. By species-level F1 score – a measurement of test accuracy that considers both precision and recall – (*F*1 *score* = ((*recall*^−1^ + *precision*^−1^) / 2) − 1**) –** PanGIA is the highest scoring tool at the 1e3 read-level and above (**Figure 5**). K-mer classifiers CLARK-S, Kaiju, Kraken2, and KrakenUniq, as well as the BWT/FM read-mapping tool Centrifuge, all perform better at the lowest read-level of 100 spiked reads per organism. The conserved marker-based classifier, Metaphlan2, struggles at the low-end read-levels but outpaces all but PanGIA and GOTTCHA at the higher spikes. Metaphlan2’s behavior is understandable in light of the higher sequencing depth required to ensure coverage on differentiating marker genes. Interestingly, all k-mer based tools (plus Centrifuge) have consistent performance across all read-levels, yet never surpass ∼ 0.81 F1 score. Since the requisite match length for k-mer classifiers (and BWT/FM-derived strings like those used by Centrifuge) tends to be shorter than full-length read mapping, classifying a signal from fewer reads is perhaps more readily achievable and their higher sensitivity at lower target signal is due to shorter sequences being required for identification. However, this characteristic becomes problematic at higher read-levels as improving true-positive detection paired with consistently low false-negative rates of read-mapping-based tools like PanGIA and GOTTCHA overcomes the consistently higher false-positive rate of k-mer (and BWT/FM) tools (**Supplementary Table 1**). Via Youden’s J score (**Supplementary Figure S1**), tested classifiers separate more distinctly. PanGIA exhibited the highest Youden’s J score across the entire titer-level range, followed by GOTTCHA (however, GOTTCHA was outperformed by Kraken2, KrakenUniq, and CLARK-S at the lowest per-target spike level (**Supplementary Figure S1, Supplementary Table 1**). Kraken2, KrakenUniq, and CLARK-S cluster closely, ranging from ∼0.49 to 0.53 Youden’s J score, with Centrifuge and Kaiju exhibiting the lowest scores across the full titer range.

**Figure 5:**
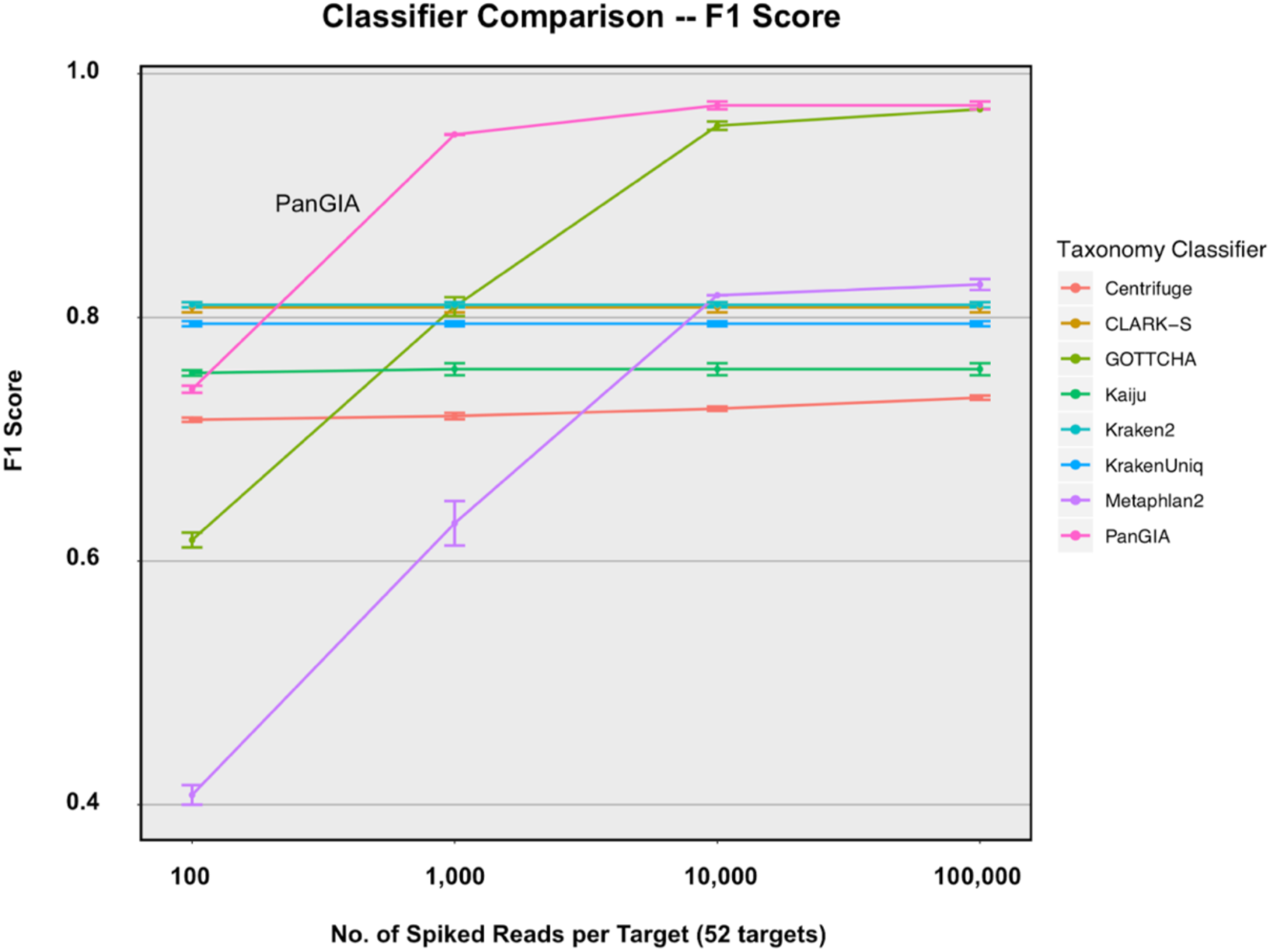
F1 Score statistic across increasing signal of 52 bacterial and viral targets generated in-silico and spiked into ‘real world’ processed forensic swab sample as background.

Tools exhibiting high false-positive-rates have reduced specificity, but the range of their misclassifications is not captured by F1 score. To help quantify this range, classification results from increasing levels of signal from individual organisms were obtained. Classifiers exhibiting flat F1 scores (**Figure 5**) attributable to low specificity (**Supplementary Table 1**) were distinctly identifiable in single-organism “off-target” analysis. When classifying an increasing Gram-negative bacterial signal of *Burkholderia mallei*, Centrifuge, Kaiju, Kraken2, and KrakenUniq (as well as CLARK-S to a lesser extent) show high numbers of reads incorrectly classified to near-neighbor species, while GOTTCHA, Metaphlan2, and PanGIA have minimal off-target detection (**Figure 6**). Repeating the analysis for a representative RNA virus (Venezuelan Equine Encephalitis virus), the same trend is observed, indicating the behavior is independent of target genome size or phylogeny. The analysis was done for a representative Gram-positive bacterium (*S. aureus*) and DNA virus (Variola virus), with similar results observed (**Supplementary Figure S2A, S2B, Supplementary Table 2**). Kaiju, Kraken2, and KrakenUniq consistently undercalled the number of *B. mallei* reads by an order of magnitude, and Kaiju continued this behavior in the *S. aureus* analysis. Some tools showed inconsistent detection across the read-abundance levels (i.e., CLARK-S and Metaphlan2). For example, CLARK-S could not identify species-specific reads accurately at the 1,000-read level, but could do so with varying degrees of recall at the 100-read level for all organisms (**Figure 6, Supplementary Figure S2**). Metaphlan2 identified species-level Variola virus signal *only* at the 10,000-read level – at the 100,000-read level, the classification was for the vaccine strain of Variola (Vaccinia virus). While *Orthopox* reads were identified across all titer-levels, the reliance on specific markers may beget inconsistencies in species-level detection of certain organisms with Metaphlan2, especially if regions of the target genome containing the markers are biased against in amplification strategies for shotgun sequencing (e.g., WGA), though this does not explain this odd behavior with *in silico* data. We considered that the CLARK-S and Metaphlan2 behavior may have been an error, but orthogonal analyses (including with re-made *in-silico* reads) confirmed the results. GOTTCHA and Metaphlan2 consistently match PanGIA’s low off-target detection, however, they also exhibit lower recovery of species-level signal in the lower titer levels due to reliance on marker (or signature)-based databases. PanGIA is the only tool that accurately classifies all four organisms to the species-level at 1,000-read abundance and above, with minimal off-target detection.

**Figure 6:**
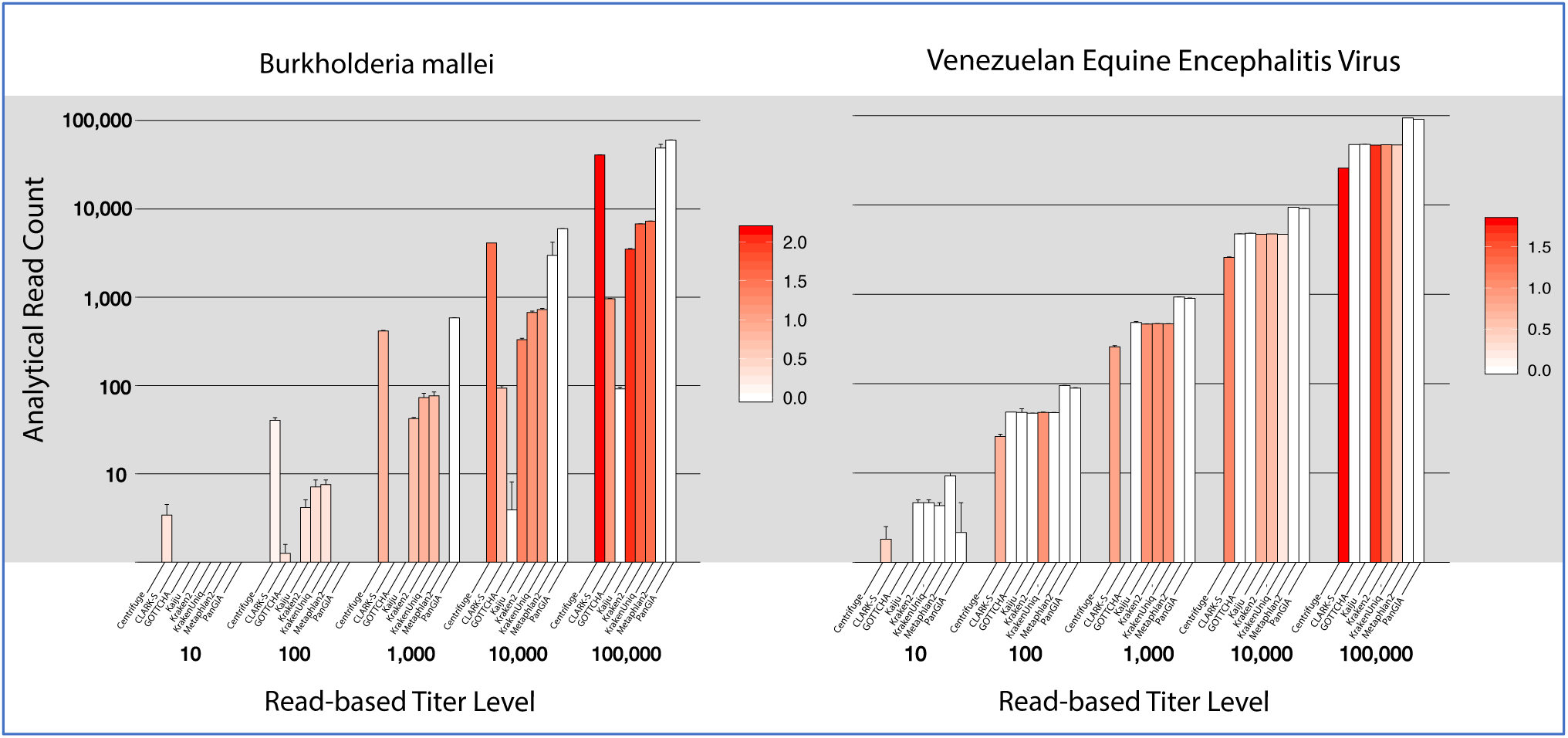
Raw target read recovery and representation of near-neighbor off-target assignments for two model organisms (Gram Negative, and RNA virus) for the 8 benchmarked taxonomy classifiers (from left to right in each grouping; Centrifuge, CLARK-S, GOTTCHA, Kaiju, Kraken2, KrakenUniq, Metaphlan2, and PanGIA) – reported at the species level. The Y-axis shows the number of reads recovered; intensity of color represents the number of off-target species-level assignments returned by the tool (i.e., LOG_10_ number of species returned, minus 1 to subtract for the on-target species).

### Interactive use of PanGIA and data visualization

We developed an interactive web-based tool, IMTV, to help visualize results and quickly parse the most relevant information from large volumes of information (**Figure 7**). On the top of the web page, a dashboard reveals overall statistics of mapped/unmapped/host reads and the high-level distribution of current selected results. The overview section also displays taxonomy profiling results in a sortable table (**Figure 7A)**. Below the table is a dot plot of observed organisms with a right-side panel of tunable parameters that users may adjust/filter to interactively view the impact on the results. An example of the impact of these parameters in ‘sorting’ raw results in to high-confidence hits and relevant, actionable information can be followed in **Supplementary Figure S3**. In Panel A of Supplementary Figure S3, the full output of an analyzed sample can be seen. In Panel B, the ‘*Pathogen Only*’ button is engaged and all taxa that are not known pathogens are dropped from the IMTV dotplot – a reduction from 60 taxa to 34 taxa. In Panel C, the ‘*Minimum Genome Coverage*’ slider is adjusted to 0.1 (10% linear coverage of each taxon’s genome), and we further reduce the displayed taxa from 34 to 23. In Panel D, the ‘*Minimum Score*’ is adjusted to 1.0 (minimum standalone confidence score), and we are left with 6 pathogens detected at extremely high confidence. This dynamic analysis capability is also shown in **Figure 8A** and **8B**. The size of the circles in the dot plot represents the relative abundance of identified organisms, with the taxonomic names displayed on the x-axis and the normalized read count RNR on the y-axis. The color of the dot indicates the confidence from low to high using a blue-yellow-red color scale. By default, PanGIA reports only those species (bacteria, archaea, viruses) that are known or suspected pathogens (**Figure 7B;** see Methods), but can display all organisms found when ‘All Taxonomies’ is selected. IMTV offers the freedom to change what these attributes represent (i.e., the display can be custom set such that size represents confidence score, color represents abundance, etc.). The taxonomic level can be toggled between ranks (from superkingdom to strain). When strain level is used, IMTV will additionally display the genome map of a strain (selected via the table, or the dot plot), with the read depth-of-coverage along the length of the selected genome. The coverage plot can also highlight previously characterized regions, including species-specific sections of the genome, antimicrobial resistance (AMR) genes, toxins or virulence markers (**Figure 7C;** see Methods). The bottom of the page displays pathogen metadata such as known location, source, host and disease (if available), as well as the major read mapping statistics of the selected strain, including rank specific read counts and identity statistics. Closer views of the IMTV results shown in Figure 7 can be found in Supplemental Figures S4-S6.

**Figure 7:**
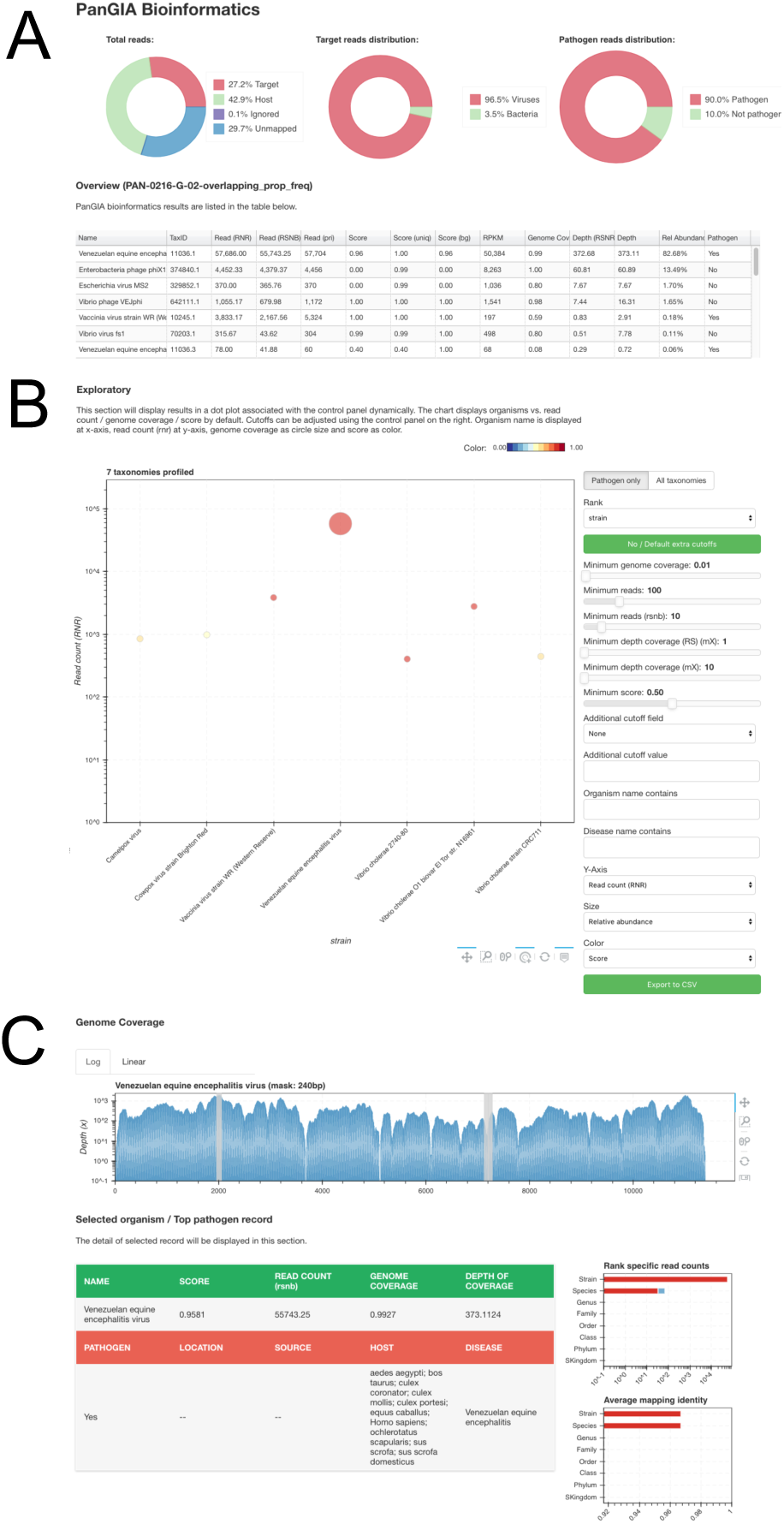
Full view of the visualization of results through IMTV. (A) The upper section of the visualization provides a dashboard of graphics with statistics concerning the categories the data was determined to contain: host, mapped, unmapped. (B) The middle section shows a dot plot with all of the pathogens identified with default parameters. (C) The lower section shows a genome coverage plot for a specific genome identified in the sample when the user selects to show strain level and clicks on the row in the table or dot in the dot plot. The gray bars indicate regions pre-identified as relevant/important marker genes (see Methods). Additionally, there is metadata provided for the selected pathogen/organism. (See Supplementary Figures S4-S6 for closer views of the IMTV results.)

**Figure 8.**
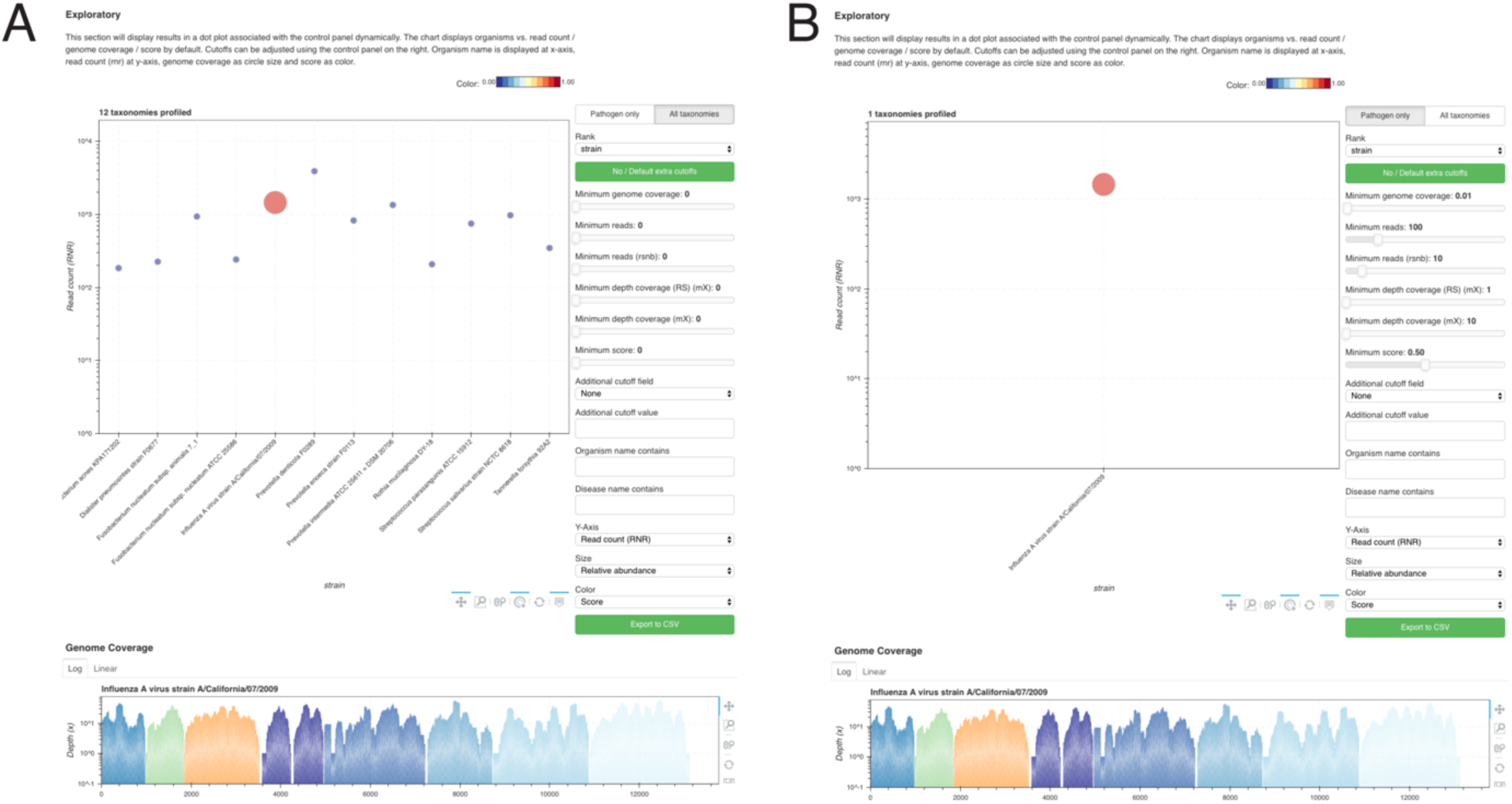
Result visualization using IMTV for a real-world clinical dataset. This is from a nasopharyngeal swab dataset where Influenza A is detected. Strain level is selected to allow the visualization of coverage of the reference genome, shown at the bottom of each of the dot plots. The genome coverage plot shows high coverage of all eight segments of the virus. (A) “All Taxonomies” was selected and all filters were dropped to zero to show all potentially identified members of the microbiome in the sample. (B) There is only one pathogen identified with the default settings which include “Pathogens only” and with a minimum confidence score of 0.5.

### Detection of pathogens in real clinical samples

After analyzing the simulated datasets, we tested how PanGIA performs on the clinical datasets. One dataset was collected from a nasopharyngeal swab from a patient with symptoms of influenza. The profiling results show ∼76% of the input reads map well to the human reference genome and only 0.2% reads mapped to the microbial references. *Influenza A* is reported as the most abundant species. At the strain level, all 8 segments of Influenza A virus strain/California/07/2009 were well covered and provide a near-complete genome (**Figure 8**). Another nasopharyngeal swab from a patient with a suspected case of pertussis was also confirmed with PanGIA. In this sample, *Bordetella pertussis* is the only pathogen species found, and strain Tohama I is the only strain identified. A human blood serum sample was also examined. Dengue virus was identified as the most abundant pathogen in this sample and is the only pathogen reported in IMTV with the default settings. (See Supplemental Figures S7-S9 for full visualization results for the clinical samples.)

We compared PanGIA to previously published taxonomy classifiers (SURPI [37] and Taxonomer [38]) using the datasets published in those manuscripts. This comparison data is in **Supplementary Table 3**. We found that PanGIA identifies all of the known etiological agent organisms identified by SURPI and Taxonomer, as well as species-level assignments for *Leptospira santarosai* not reported by SURPI at the time of publication. Wilson and colleagues used SURPI to identify patients with neuroleptospirosis infections [39]. Samples are identified by their SRA ID number. Samples SRR1145846, SRR1145844, SRR1145847, and SRR1145845 are associated with cerebrospinal fluid (CSF) from a *Leptospira santarosai* infected patient, DNAse-treated CSF from the same patient, serum from the same patient, and DNAse-treated serum from an unrelated patient, respectively. PanGIA recapitulates the strongest signal for the etiological agent in the untreated CSF sample, seen by SURPI as 475 reads reported to the genus level for *Leptospira*, and reported by PanGIA with an RNR value of 3,112 and a standalone confidence score value of 0.22 for a species-level call of *Leptospira santarosai.* Progressively lower genus-level *Leptospira* signal via SURPI is seen in the other sample extraction procedures (e.g., with cyanase treatment), sample matrices (e.g., serum), and an unrelated patient, and this is also observed in the species-level *Leptospira santarosai* signal via PanGIA’s RNR and standalone confidence scores. Wilson and colleagues report the closest species-level assignment for these samples via SURPI analysis was *Leptospira borgpetersenii*, and that a treatment decision for neuroleptospirosis was made from this data. Follow-up analyses confirmed that the true etiological species was *L. santarosai*. While this particular instance allowed for effective treatment decisions without an exact species identification, not all infections can be controlled from that level of granularity. We are confident that *L. santarosai* would have been detected by SURPI had it been present in SURPI’s database at the time of the analysis by Wilson and colleagues. We do not highlight this instance to distinguish PanGIA’s taxonomic classification ability, but as an example of the need for rapid database flexibility in metagenomics taxonomy classifiers if they are to be optimally deployed in clinical settings.

### PanGIA Database Flexibility

A common obstacle to effective use of metagenomics as a clinical tool is that of default databases from available taxonomy classifiers not containing a reference genome for a particular pathogen-of-interest. This would render the tool useless to practitioners concerned with the pathogen-of-interest, but who are not trained in how to develop custom database update scripts for open-source classifiers. To ameliorate this, we have implemented a ‘Database Update’ feature within PanGIA that is easily executed from within the GUI. If a user has a reference genome for an organism in FASTA format, this genome can be uploaded to PanGIA’s database during analytical run setup (**Supplementary Figure S10**). The user simply names the new database entry, selects the FASTA file from a standard directory navigation tool, and labels it as ‘*Host*’ or ‘*Target*’. Upon starting the analytical run, PanGIA will create a BWA index for the FASTA file reference genome in PanGIA’s database directory before continuing with taxonomy classification. If the FASTA reference has been labeled as ‘*Target*’, then any sequence reads mapping to the reference will be included in PanGIA’s results. If the FASTA reference has been labeled as ‘*Host*’, then any sequence reads mapping to the reference will be excluded from the results. The practical utility of this feature is demonstrated in **Supplementary Table 3**, containing the comparison of PanGIA results to those of SURPI and Taxonomer. The primary results for the initial PanGIA comparison of the sample with SRA ID # SRR1106119 are ‘NA’, or ‘not applicable’, due to an absence of PanGIA classification of those reads since a reference genome for the etiological pathogen (Bas-Congo Virus) was not part of PanGIA’s native database. However, upon re-analysis with simultaneous database update to include Bas-Congo Virus as a *target* organism, indicated in parentheses to the right of the original results in **Supplementary Table 3**, we see that PanGIA then detects the Bas-Congo Virus with an RNR value of 62,323. The detection is missing a standalone confidence score because database additions through the GUI *only* index the reference genome and make it available for mapping, which is manageable on the CPU/RAM configuration of typical commodity hardware. The process does *not* include the re-calculation of the uniqueness indices necessary for standalone confidence scoring, which requires much more CPU/RAM capability than is available on standard commodity computational equipment. If a user has access to high-performance computing environments, the process for updating their own uniqueness indices can be made available.

### PanGIA Executive Summary Report (ESR)

In order to provide concise results to end-users, an Executive Summary Report of PanGIA analytical results is populated in a fixed format (PDF), incorporating all ‘default’ settings (preventing unintentional user adjustment of results) and conveying the most critical results clearly and succinctly with minimal jargon. In clinical settings, the most actionable end-user information, ultimately, are the pathogens that were detected (if any) and the abundance and confidence scoring metrics for those hits. These are displayed in large clear lettering, using font coloring and bolding to draw the eye to the pathogen information. If no pathogen is detected, this will be displayed in similarly bold fashion. Following pathogen detection information, the most actionable information is the reliability of the results. Deviations from acceptable values at each of the built-in quality check-points would point to contamination or otherwise erroneous results. Quality control values are displayed at these checkpoints in a timeline fashion (split between the three major workflow junctions; sample preparation, sequencing, and bioinformatics), such that should there be a deviation from nominal values, the step in the workflow where the problem occurred can be quickly determined. An example ESR is shown in **Supplementary File 2.**

## DISCUSSION

Metagenomics sequencing has emerged as a promising new technology for pathogen detection and agnostic biosurveillance, particularly when the etiological agent of concern is not known. The challenge in metagenomics for any pathogen identification algorithm, and more generally for any taxonomy classification algorithm, has always been the tradeoff between sensitivity and specificity. For example, kmer-based methods are frequently cited in literature due to speed and high sensitivity, although based on the analysis presented here and elsewhere, they often exhibit a higher false positive rate. On the other hand, marker-based methods have very high specificity largely owing to curated gene lists or biomarker signatures, but while they do have the capability of reaching similar levels of sensitivity as other methods, in some instances, sensitivity can be compromised. Most alignment-based methods tolerate sequence variants well and provide contextual, location-specific information, such as depth of coverage and observed variance, but typically have longer analysis wall-clock times and are associated with higher false positives due to conserved regions shared among closely related genomes. PanGIA stands as a novel alignment-based taxonomy classification algorithm with significantly-improved sensitivity by addressing the shortcomings of these other methods. By analyzing how reads map at each taxonomic level to pre-computed taxonomy-based unique signatures, we derive a confidence metric using the unique signatures as our prior knowledge to help determine whether that particular taxon is truly in the sample.

Broad adoption of metagenomics for pathogen identification has primarily been hindered by the computational burden, the complexity of data analysis implementations, and the difficulty in automating interpretation and reporting within a routine clinical or public-health setting. In this work, we present a novel algorithm, PanGIA, for pathogen detection from metagenomics datasets that can be deployed on commodity computational hardware (e.g., standard desktop or laptop computers), can be executed from an intuitive and *dynamic* graphical user interface by non-expert end-users, and automatically outputs succinct data products designed to be rapidly interpreted by decision-makers. By integrating two separate confidence score metrics (i.e., one based on sequence-derived phylogenetic uniqueness and another based on abundance-derived information across operational control samples), PanGIA provides end-users without formal bioinformatics training a novel ability to quickly transfer the most actionable genomics-based information where it needs to be sent in mission-critical or clinically-relevant timeframes.

PanGIA demonstrates superior overall performance (determined via analytical sensitivity- and specificity-based metrics, as well as ‘off-target’ analysis) when compared to several other leading metagenomics taxonomy classifiers. In addition to confidence scores and balanced sensitivity/specificity, our tool integrates key features that layer additional supporting information to each taxonomic classification; 1) strain-level depth-of-coverage plots for each detected taxa (useful for determining how sequence reads are distributed across a target genome, particularly in segmented viruses or between bacterial chromosomes and plasmids), and 2) taxa-specific and phenotype-specific (e.g., AMR, toxicity, and virulence) nucleotide-level markers associated with each strain are displayed at their correct coordinate position on the genome coverage plot (useful for confirming strain/species level assignment, as well as clinical and/or public-health-related phenotypes-of-concern). While default settings have been optimized for balanced sensitivity and specificity across ‘generalized’ sample types, PanGIA is not a static tool. Substantial exploratory capability has been built into the platform, such that users can adjust both pre-run analytical settings and post-run result metric ‘cut-off’ values to widen or narrow the analytical scope, as appropriate. For example, by clicking the ‘Pathogen Discovery’ button in the run-time parameter settings module, default values for minimum alignment length, minimum alignment score, minimum linear coverage, minimum depth-of-coverage, minimum number of reads, and minimum confidence score are relaxed to ‘cast a wider net’ with lower required fidelity to the known reference genomes in the database. Several of these metrics can be dynamically reset on the ‘back-end’, in the IMTV module, such that displayed results adjust to updated cut-offs in real-time.

## Conclusions

PanGIA was developed to address long-standing roadblocks for the use of metagenomics technology in routine biosurveillance and clinical contexts, primarily by genomics and bioinformatics novices. By integrating novel confidence metrics, on-the-fly database flexibility, dynamic data exploration capability, and other standard sequencing-derived information into an intuitive GUI, it is our hope that the ‘barrier-to-entry’ of metagenomics in applied settings will be lowered substantially. Currently, PanGIA can analyze sequencing reads generated from the Illumina platform. Support for other sequencing technologies, e.g., Nanopore and PacBio, are currently under active development and testing, and will be included in the next major release.

## ACKNOWLEDGEMENTS

The authors would like to thank the Defense Threat Reduction Agency for funding this work.

## SUPPLEMENTARY MATERIALS

**Supplementary Figure S1:**
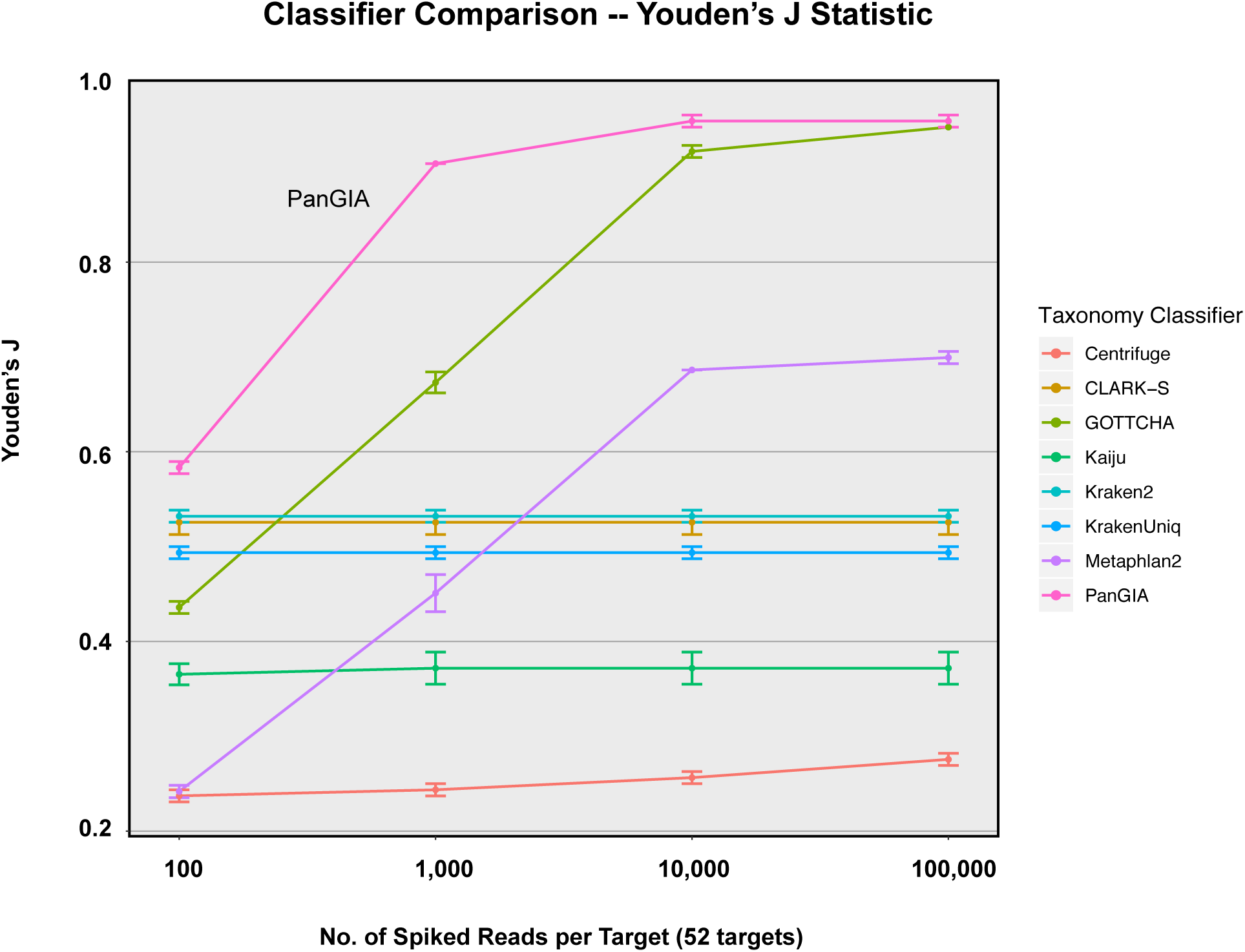
Youden’s J statistic across increasing signal of 52 bacterial and viral targets generated in-silico and spiked into ‘real world’ processed forensic swab sample as background. True positive (TP) is defined here as detection of each target (any number of reads) in samples that they were known to be spiked into. True negative (TN) is defined here as lack of detection of each target that was known to be absent in the forensic swab sample background. False positive (FP) is defined as detection of a spiked target in the forensic swab background alone. False negative (FN) is defined as lack of detection of each target that was known to be spiked into the analytical samples. Youden’s J = ((TP)/(TP+FN)) + ((TN)/(TN+FP))-1.

**Supplementary Figure S2:**
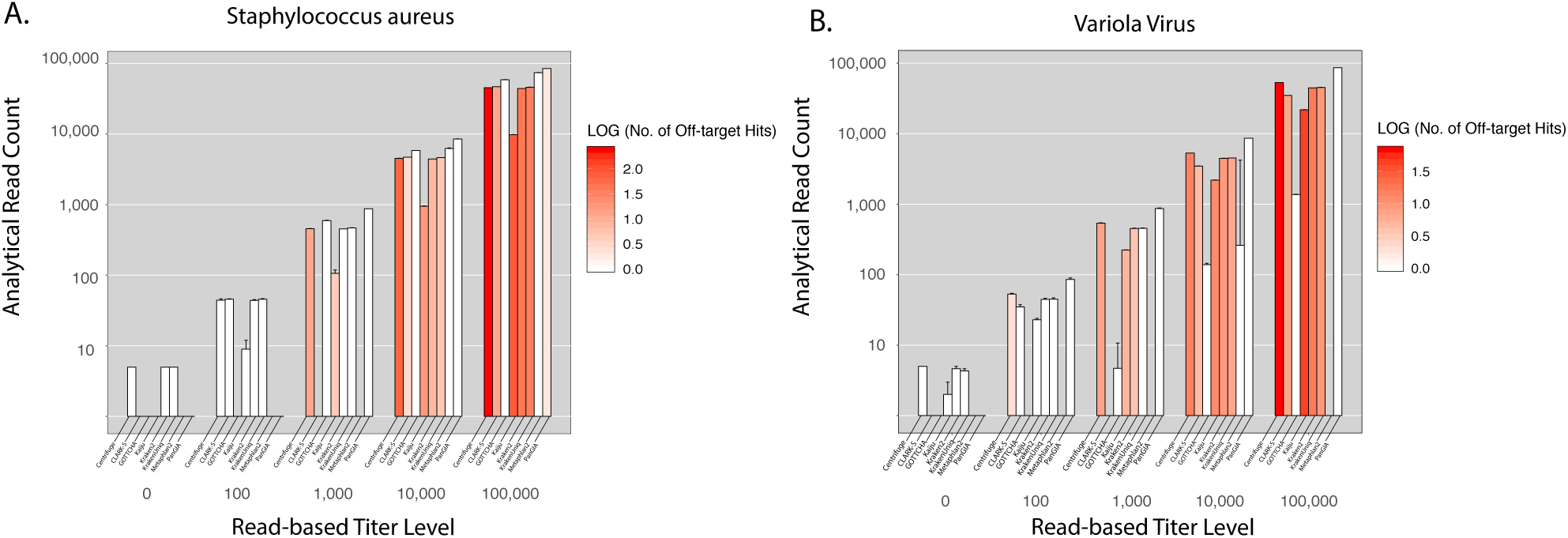
Raw target read recovery and representation of near-neighbor off-target assignments for a Gram + (S. aureus, **A**) and DNA virus (Variola virus, **B**) model organisms across the 8 benchmarked metagenomic read classifiers, reported at the species level. Raw reads were generated in-silico from each species target at increasing mock titer levels (10; 100; 1,000; 10,000; 100,000) and classified with each tool (default settings -- listed in Supplementary material). This was conducted to model the performance of each tool in an “ideal” classification scenario (i.e., if we give 100,000 reads of species-X to tool Y, that tool should return 100,000 reads of species-X, and nothing else). The Y-axis shows the number of reads recovered and intensity of color associated with each tool at each titer level represents the number of off-target species-level assignments returned by the tool (i.e., number of species returned minus 1 (LOG_10_)).

**Supplementary Figure S3:**
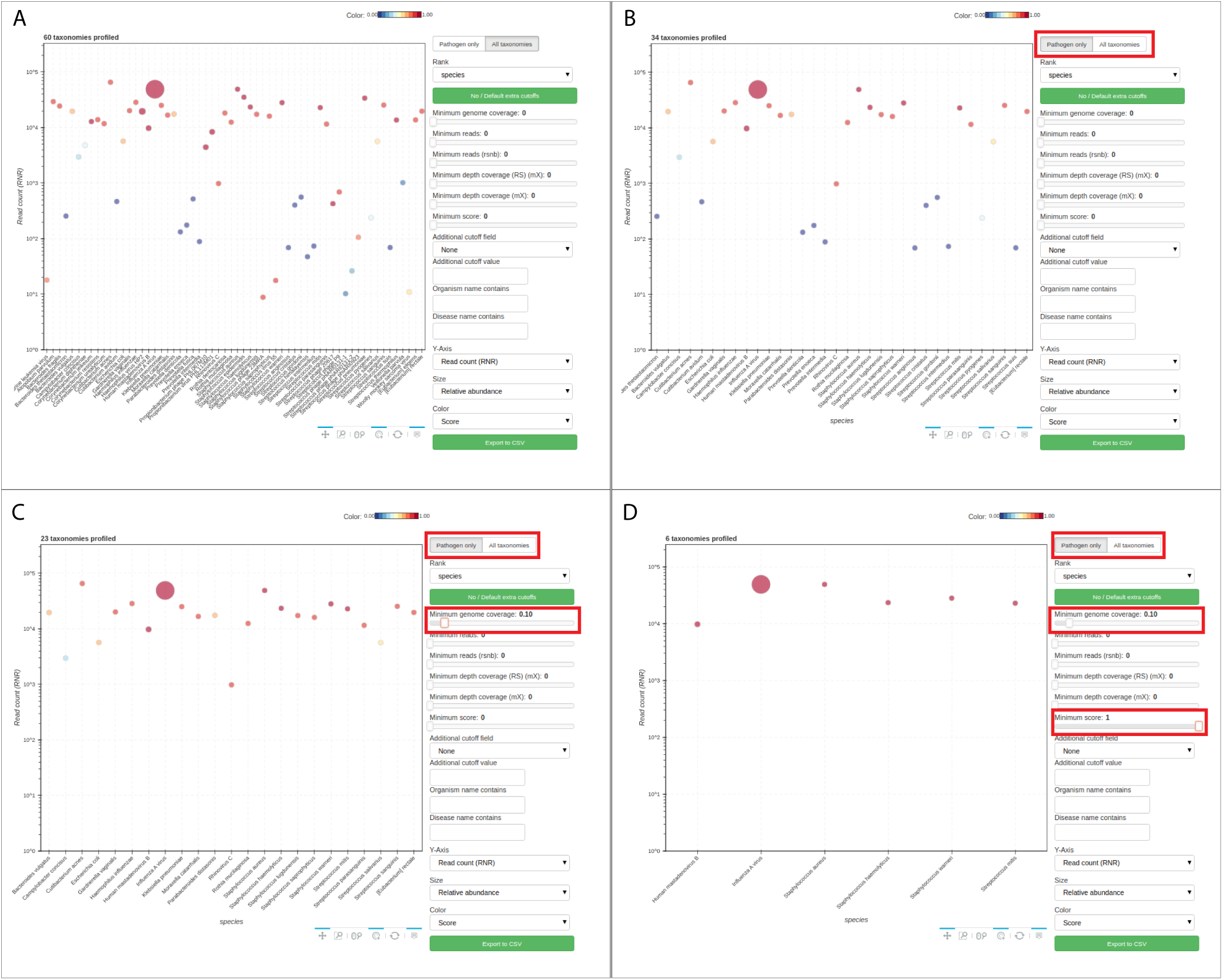
Use of real-time ‘toggles’ in PanGIA’s IMTV to dynamically sort raw results to only pathogens detected at high-confidence in a matter of seconds.

**Supplementary Figure S4:**
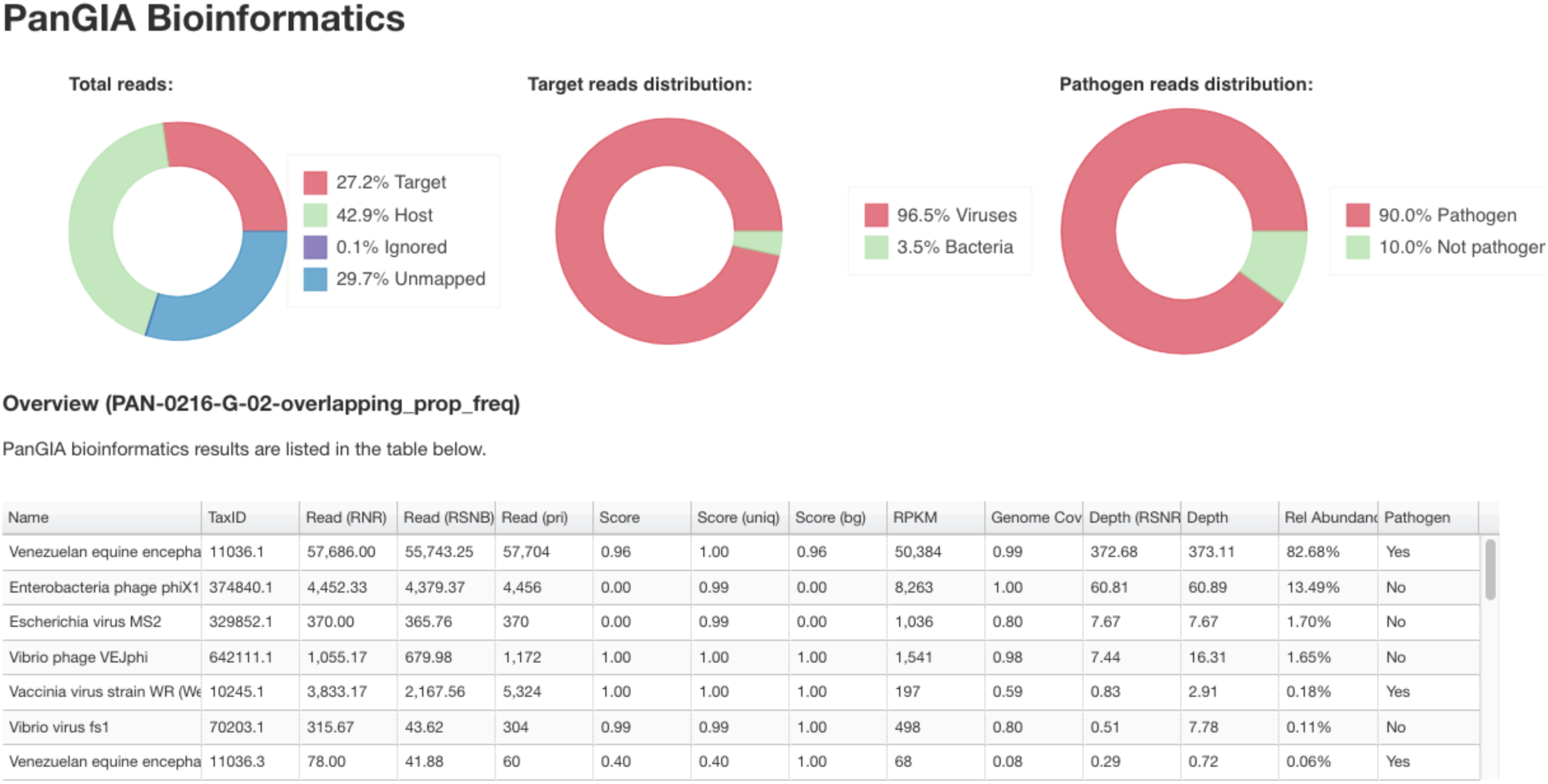
A closer view of the upper section of results visualization of results in IMTV. The dashboard provides graphics with statistics of what was identified in the sample. This sample was one of the spiked sets described earlier in the main text.

**Supplementary Figure S5:**
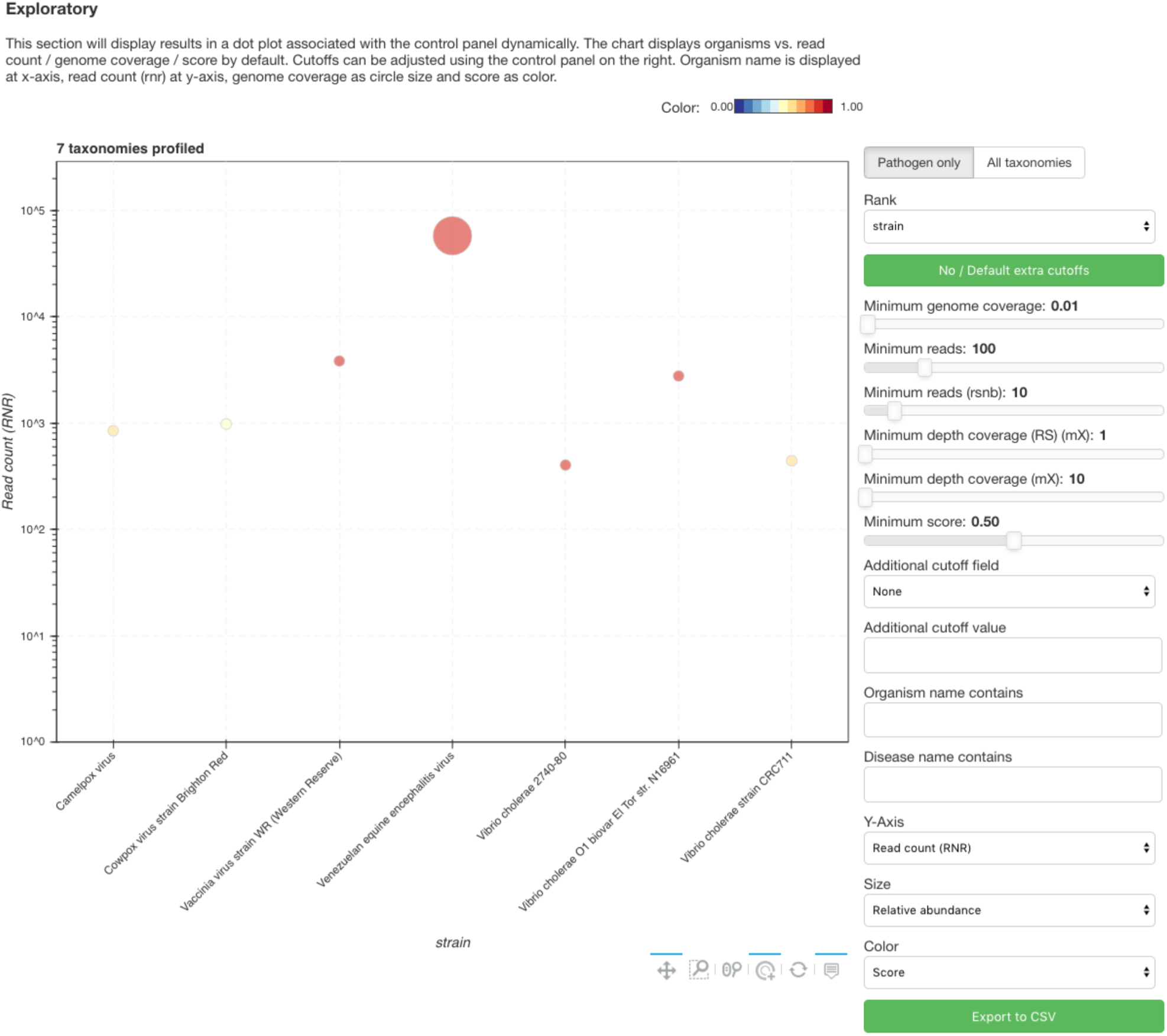
A closer view of the middle section of results visualization of results in IMTV. The dot plot shows pathogens identified with default parameters.

**Supplementary Figure S6:**
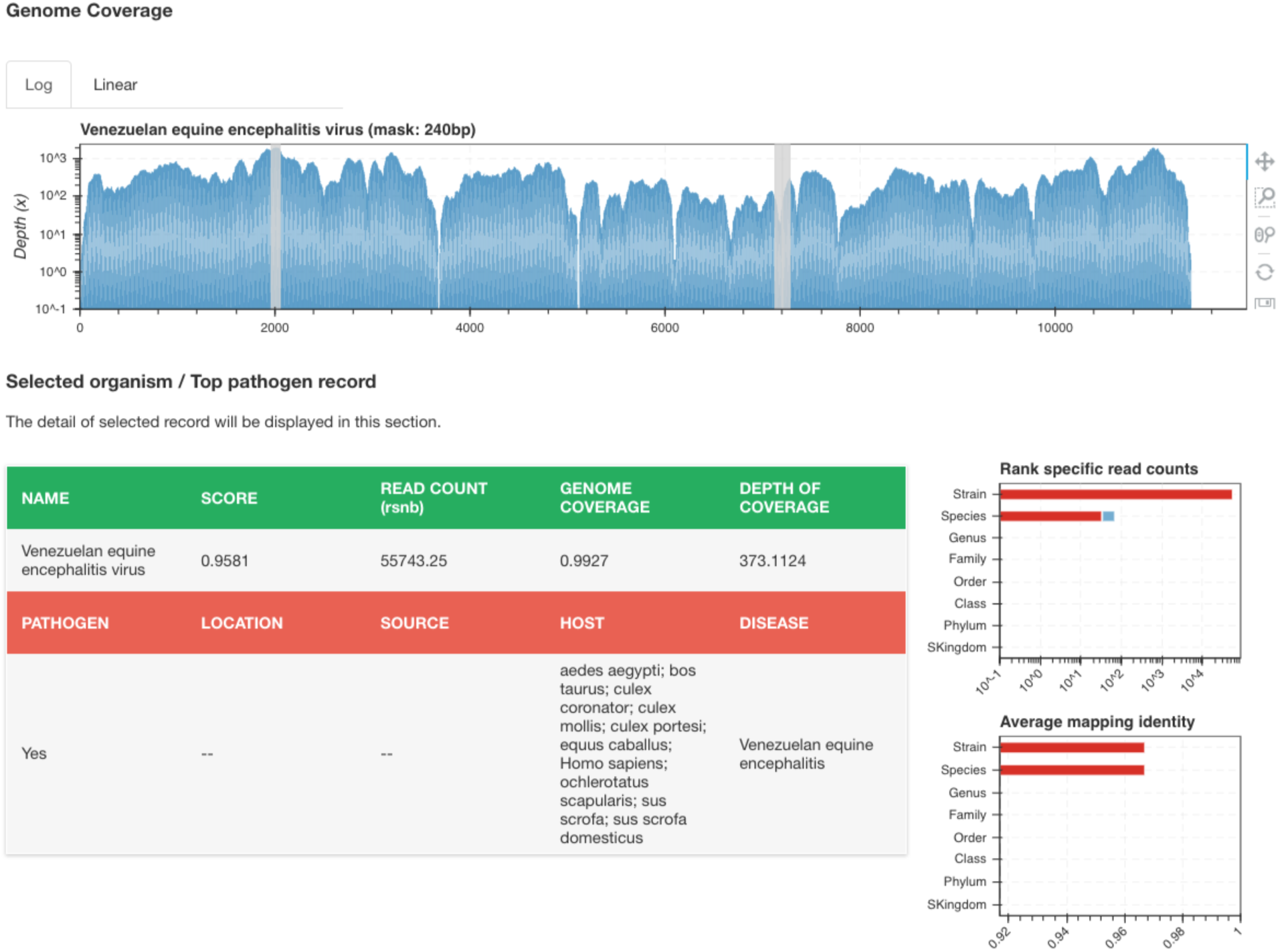
A closer view of the lower section of results visualization of results in IMTV. When strain level and a particular pathogen are selected from the dot plot or the table above, a genome coverage plot of the data mapped to the reference is shown. The gray bars indicate regions mapped to in the negative control indicating potential contamination from VEEV amplicons. Metadata concerning the pathogen are also provided along with histograms showing number of reads mapping at specific taxanomic ranks.

***Supplementary Figure S7:** Result visualization using IMTV for real-world datasets. Each box shows the visualization of results for the clinical samples at strain level with default filter settings. Strain level allows the visualization of coverage of the reference genome, shown at the bottom of each figure. In each of these examples, there is only one pathogen identified with the default setting with a minimum confidence score of 0.5. A nasopharyngeal swab dataset where Influenza A is detected. The genome coverage plot shows high coverage of all eight segments of the virus.*

***Supplementary Figure S8.** Bordetella pertussis was detected with high coverage of the entire genome with only a few islands not being represented in the sequence data.*

***Supplementary Figure S9.** Results of a serum sample detects Dengue virus. Abundant but incomplete genome coverage of the reference indicates the sample contains a different strain of Dengue virus than the reference provided in the database.*

**Supplementary Figure 10:**
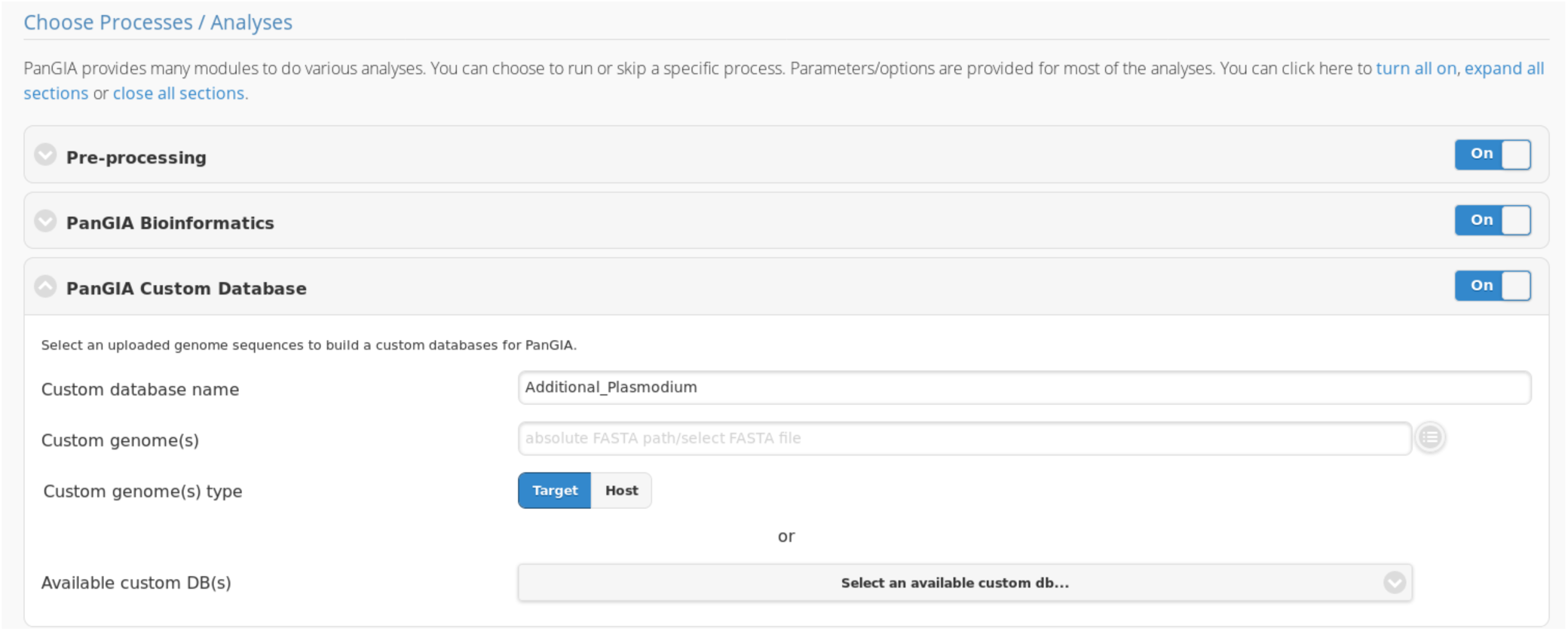
Database update/customization within the PanGIA GUI.

***Supplementary File 1 (attachment)***: Command-line submission scripts for each tool benchmarked against PanGIA, as well as the list of organisms included in *in-silico*-spiked benchmarking datasets.

***Supplementary File 2 (attachment)***: *PanGIA Executive Summary Reports provide actionable information. The summary report gives a high-level overview of the sample, pathogens identified with confidence level, and a pass or fail call for Quality Assurance for each of the following steps: sample preparation, sequencing, and bioinformatic analysis.*

***Supplementary Table 1 (attachment)***: *Full benchmarking data of PanGIA against other leading taxonomy classifiers*

***Supplementary Table 2 (attachment)***: *Full benchmarking data of near-neighbor off-target analysis for PanGIA against other leading taxonomy classifiers*

***Supplementary Table 3 (attachment)***: *Comparison of PanGIA performance on benchmarking data from the SURPI and Taxonomer publications.*

