## Supplementary File 1 for "PanGIA: A Metagenomics Analytical Framework for Routine Biosurveillance and Clinical Pathogen Detection"

**#*Awk* and dwgsim commands for *in-silico* sequence read generation**

**#Example of sub-sampling reads from environmental background sequencing data to generate a 10M read 'background' dataset**

```
paste env_background_R1.fastq env_background_R2.fastq | awk '{
printf("%s",$0); n++; if(n%4==0) { printf("\n");} else { printf("\t");} }' |
awk -v k=10000000 'BEGIN{srand(systime() +
PROCINFO["pid"]);}{s=x++<k?x-
1:int(rand()*x);if(s<k)R[s]=$0}END{for(i in R)print R[i]}' | awk -F"\t"
'{print $1"\n"$3"\n"$5"\n"$7 >
"env_background_10000000_rep1_R1.fastq";print
$2"\n"$4"\n"$6"\n"$8 > "env_background_10000000_rep1_R2.fastq"}'
```

**#Examples of *dwgsim* commands to generate 5,000 paired-end 2x75bp from virus and bacteria genomes**

```
for i in /viruses/*fa; do name=$(basename $i .fa); dwgsim -c 0 -S 2 -e
0.001-0.01 -E 0.001-0.01 -d 500 -s 10 -1 75 -2 75 -n 0 -N 5000 $i >
/$name.fastq; done
```

```
for i in /bacteria/*fa; do name=$(basename $i .fa); dwgsim -c 0 -S 2 -e
0.001-0.01 -E 0.001-0.01 -d 500 -s 10 -1 75 -2 75 -n 0 -N 5000 $i >
$name.fastq; done
```

#Representative Metagenomic Community Dataset
#Analysis Scripts for Each Tested Classifier

**#PanGIA**

```
DATALOC="/ /";
RESULTSLOC="/ /";
TOOLLOC="/ /PanGIA";
TOOLDATABASE="/ /PanGIA/database/";

FASTQ=($(ls $DATALOC/*fastq));

for ((i=0; i<${#FASTQ[@]};i++));

do
NAME=$(echo ${FASTQ[$i]} | sed 's/.fastq//');
NAME=$(echo $NAME | awk -F/ '{print $NF}');
echo "Sample $NAME: Processing ${FASTQ[$i]}     ${FASTQ[$i+1]}";

    /usr/bin/python3.6 $TOOLLOC/pangia.py \
        -d $TOOLDATABASE/*.fa \
        -i ${FASTQ[$i]} ${FASTQ[$i+1]} \
        -t 32 \
        -ams 60 \
        -asl 40 \
        -td /tmp/  \
        -o $RESULTSLOC/$NAME \

        i=$((i+1))

done;
```

#### #Centrifuge

```
set -xeu

DATALOC="/ /";
TOOLLOC="/ /centrifuge";
RESULTSLOC="// ";

FASTQS=($(ls $DATALOC/ *.fastq));

echo $FASTQS

for ((i=0; i<${#FASTQS[@]};i++));
do

   NAME=$(echo ${FASTQS[$i]} | sed 's/.fastq//');
   NAME=$(echo $NAME | awk -F/ '{print $NF}');
   NAME=$(echo $NAME | sed 's/_QCB_R.//');

$TOOLLOC/centrifuge -x $TOOLLOC/indices/nt -1 ${FASTQS[$i]} -2 ${FASTQS[$i+1]} -S
$RESULTSLOC/${NAME}_hits.tsv -p 12 --report-file $RESULTSLOC/${NAME}_report.tsv

   i=$((i+1));

done;
```

### #CLARK-S

```
set -xeu

DATALOC="/ /";
RESULTSLOC="/ /";

FASTQ=($(ls $DATALOC/*.fastq));

for ((i=0; i<${#FASTQ[@]};i++));
do
   NAME=$(echo ${FASTQ[$i]} | sed 's/.fastq//');
   NAME=$(echo $NAME | awk -F/ '{print $NF}');
   NAME=$(echo $NAME | sed 's/_QCB_R.//');
   echo "Sample $NAME";

 bash /home/src/CLARKSCV1.2.3/set_targets.sh /home/src/CLARKSCV1.2.3/DIR_DB2
bacteria viruses --species

bash /home/src/CLARKSCV1.2.3/classify_metagenome.sh -P ${FASTQ[$i]} ${FASTQ[$i+1]}
-R ${RESULTSLOC}/{$NAME}_results --spaced -n 32

bash /home/src/CLARKSCV1.2.3/estimate_abundance.sh -F
${RESULTSLOC}/{$NAME}_results.csv -D /home/src/CLARKSCV1.2.3/DIR_DB2  >
${RESULTSLOC}/{$NAME}_taxa.tsv

   i=$((i+1));

done;
```

**#GOTTCHA**

```
set -xeu

DATALOC="/ /";
RESULTSLOC="/ /";

GOTTCHAdbBACTERIA="/home/src/gottcha/database/GOTTCHA_BACTERIA_c4937_k24_u30_xHUMAN3x.species";
GOTTCHAdbVIRUSES="/home/src/gottcha/database/GOTTCHA_VIRUSES_c5900_k24_u30_xHUMAN3x.species";

FASTQ=($(ls $DATALOC/*.fastq));

for ((i=0; i<${#FASTQ[@]};i++));
do
    NAME=$(echo ${FASTQ[$i]} | sed 's/.fastq//');
    NAME=$(echo $NAME | awk -F/ '{print $NF}');
    NAME=$(echo $NAME | sed 's/_QCB_R.//');
    echo "Sample $NAME";

    /home/bin/gottcha        \
        --threads 16        \
        --outdir $RESULTSLOC \
        --input ${FASTQ[$i]} \
        --prefix $NAME.BACTERIA.1 \
        --database   $GOTTCHAdbBACTERIA

    /home/bin/gottcha        \
        --threads 16        \
        --outdir $RESULTSLOC \
        --input ${FASTQ[$i]} \
        --prefix $NAME.VIRUSES.1\
        --database   $GOTTCHAdbVIRUSES

sed '2,$!d' $RESULTSLOC/$NAME.VIRUSES.1.gottcha.tsv > $RESULTSLOC/$NAME.VIRUSES.1.gottcha.part.tsv
```

```
cat $RESULTSLOC/$NAME.BACTERIA.1.gottcha.tsv
$RESULTSLOC/$NAME.VIRUSES.1.gottcha.part.tsv >
$RESULTSLOC/$NAME.BACandVIR.1.gottcha.tsv


   /home/bin/gottcha        \
      --threads 16        \
      --outdir $RESULTSLOC \
      --input ${FASTQ[$i+1]} \
      --prefix $NAME.BACTERIA.2 \
      --database   $GOTTCHAdbBACTERIA

   /home/bin/gottcha        \
      --threads 16        \
      --outdir $RESULTSLOC \
      --input ${FASTQ[$i+1]} \
      --prefix $NAME.VIRUSES.2\
      --database   $GOTTCHAdbVIRUSES

sed '2,$!d' $RESULTSLOC/$NAME.VIRUSES.2.gottcha.tsv >
$RESULTSLOC/$NAME.VIRUSES.2.gottcha.part.tsv

cat $RESULTSLOC/$NAME.BACTERIA.2.gottcha.tsv
$RESULTSLOC/$NAME.VIRUSES.2.gottcha.part.tsv >
$RESULTSLOC/$NAME.BACandVIR.2.gottcha.tsv


sed '2,$!d' $RESULTSLOC/$NAME.BACandVIR.2.gottcha.tsv >
$RESULTSLOC/$NAME.BACandVIR.2.gottcha.part.tsv

cat $RESULTSLOC/$NAME.BACandVIR.1.gottcha.tsv
$RESULTSLOC/$NAME.BACandVIR.2.gottcha.part.tsv >
$RESULTSLOC/$NAME.BACandVIR.gottcha.tsv

   i=$((i+1));

done;
```

### #Kaiju

```
set -xeu

DATALOC="/ /";
TOOLLOC="// ";
RESULTSLOC="/ /";

FASTQ=($(ls $DATALOC/*.fastq));

for ((i=0; i<${#FASTQ[@]};i++));
do
	NAME=$(echo ${FASTQ[$i]} | sed 's/.fastq//');
	NAME=$(echo $NAME | awk -F/ '{print $NF}');
	NAME=$(echo $NAME | sed 's/_QCB_R.//');
	echo "Sample $NAME: Processing ${FASTQ[$i]}    ${FASTQ[$i+1]}";

kaiju -t $TOOLLOC/kaijudb/nodes.dmp -f $TOOLLOC/kaijudb/kaiju_db.fmi -i ${FASTQ[$i]} -
j ${FASTQ[$i+1]} -o $RESULTSLOC/${NAME}_kaiju.out -v -z 16

addTaxonNames -t $TOOLLOC/kaijudb/nodes.dmp -n $TOOLLOC/kaijudb/names.dmp -i
$RESULTSLOC/${NAME}_kaiju.out -o -i $RESULTSLOC/${NAME}_kaiju-names.out

kaijuReport -t $TOOLLOC/kaijudb/nodes.dmp -n $TOOLLOC/kaijudb/names.dmp -i
$RESULTSLOC/${NAME}_kaiju.out -r species -o $RESULTSLOC/${NAME}_kaiju-
names.out.summary

	i=$((i+1));

done;
```

#### #Kraken2

```
set -xeu

DATALOC="/ /";
RESULTSLOC="/ /";

FASTQ=($(ls $DATALOC/*.fastq));

for ((i=0; i<${#FASTQ[@]};i++));
do
    NAME=$(echo ${FASTQ[$i]} | sed 's/.fastq//');
    NAME=$(echo $NAME | awk -F/ '{print $NF}');
    NAME=$(echo $NAME | sed 's/_QCB_R.//');
    echo "Sample $NAME";

    /home/bin/kraken2 \
        --paired \
        --db /home/src/kraken2-2.0.7-beta_102518/full/  \
        --threads 32 \
        --classified-out $RESULTSLOC/${NAME}-class\#.fastq \
        --unclassified-out $RESULTSLOC/${NAME}-unclass\#.fastq  \
        --output $RESULTSLOC/${NAME}-krakenout.txt  \
        --report $RESULTSLOC/${NAME}.report.txt \
        ${FASTQ[$i]} ${FASTQ[$i+1]};


    i=$((i+1));

done;
```

### #KrakenUniq

```
set -xeu

DATALOC="/ /";
RESULTSLOC="/ /";
DBDIR="/ /";

/home/bin/krakenuniq/krakenuniq --db $DBDIR --preload --threads 32

FASTQ=($(ls $DATALOC/*.fastq));

for ((i=0; i<${#FASTQ[@]};i++));
do
   NAME=$(echo ${FASTQ[$i]} | sed 's/.fastq//');
   NAME=$(echo $NAME | awk -F/ '{print $NF}');
   NAME=$(echo $NAME | sed 's/_QCB_R.//');
   echo "Sample $NAME";

   /home/bin/krakenuniq/krakenuniq \
      --paired \
      --check-names  \
      --db $DBDIR  \
      --threads 32 \
      --classified-out $RESULTSLOC/${NAME}-class\#.fastq \
      --unclassified-out $RESULTSLOC/${NAME}-unclass\#.fastq  \
      --output $RESULTSLOC/${NAME}-krakenuniqout.txt  \
      --report $RESULTSLOC/${NAME}.report.txt \
      ${FASTQ[$i]} ${FASTQ[$i+1]};


   i=$((i+1));

done;
```

#### #Metaphlan2

```
set -xeu

EXEC=/home/bin;
mpa_dir=/home/src/metaphlan2;
DATALOC="// ";
RESULTSLOC="/ /";
TEMPDIR="/ /";

FASTQS=($(ls $DATALOC/bac_vir_ROC*.fastq));

for ((i=0; i<${#FASTQS[@]};i++));
do
   NAME=$(echo ${FASTQS[$i]} | sed 's/.fastq//');
   NAME=$(echo $NAME | awk -F/ '{print $NF}');
   NAME=$(echo $NAME | sed 's/_QCB//');
   echo "Sample $NAME";

   /usr/bin/python2.7 $mpa_dir/metaphlan2.py \
      ${FASTQS[$i]} \
      --input_type multifastq \
      --nproc 32 \
      --mpa_pkl ${mpa_dir}/db_v20/mpa_v20_m200.pkl \
      --bowtie2db ${mpa_dir}/db_v20/ \
      -t rel_ab_w_read_stats \
      --bowtie2out ${RESULTSLOC}/${NAME}.bowtie2out.txt >
${RESULTSLOC}/${NAME}_profile.txt


done;
```

```
/usr/bin/python2.7 $mpa_dir/utils/merge_metaphlan_tables.py
${RESULTSLOC}/bac_vir_ROC*_profile.txt >
${RESULTSLOC}/${NAME}_merged_abundance_table.txt

FASTQB=($(ls $DATALOC/env*.fastq));

for ((i=0; i<${#FASTQB[@]};i++));
do
   NAME=$(echo ${FASTQB[$i]} | sed 's/.fastq//');
   NAME=$(echo $NAME | awk -F/ '{print $NF}');
   NAME=$(echo $NAME | sed 's/_QCB//');
   echo "Sample $NAME";

   /usr/bin/python2.7 $mpa_dir/metaphlan2.py \
      ${FASTQB[$i]} \
      --input_type multifastq \
      --nproc 32 \
      --mpa_pkl ${mpa_dir}/db_v20/mpa_v20_m200.pkl \
      --bowtie2db ${mpa_dir}/db_v20/ \
      -t rel_ab_w_read_stats \
      --bowtie2out ${RESULTSLOC}/${NAME}.bowtie2out.txt >
${RESULTSLOC}/${NAME}_profile.txt


done;

/usr/bin/python2.7 $mpa_dir/utils/merge_metaphlan_tables.py
${RESULTSLOC}/env*_profile.txt >
${RESULTSLOC}/${NAME}_merged_abundance_table.txt
```

### #List of 52 Organisms *in-silico* spiked into soil background metagenomics data

Vibrio cholerae
Yersinia pestis
Brucella melitensis
Bacillus anthracis
Burkholderia mallei
Francisella tularensis
Xanthomonas oryzae
Deinococcus geothermalis
Streptococcus pneumoniae
Bifidobacterium adolescentis
Bacteroides vulgatus
Clostridium beijerinckii
Acinetobacter baumannii
Bacillus cereus
Borrelia burgdorferi
Clostridium botulinum
Rickettsia rickettsia
Helicobacter pylori
Pseudomonas aeruginosa
Staphylococcus haemolyticus
Bacillus thuringiensis
Burkholderia pseudomallei
Human gammaherpesvirus 4
Zika virus
Gammapapillomavirus 12
Megavirus chiliensis
Tokyovirus A1
Rickettsia prowazekii
Streptococcus pyogenes
Influenza A virus
Japanese encephalitis virus
Venezuelan equine encephalitis virus
Dengue virus
Marburg Marburgvirus
Variola virus
Influenza B virus
Zaire ebolavirus
Nipah henipavirus
Cowpox virus
Enterovirus G
Hepacivirus C
Chikungunya virus
Colorado tick fever virus
Lassa mammarenavirus
Enterovirus G
Omsk hemorrhagic fever virus
Guanarito mammarenavirus
Hantaan orthohantavirus
Crimean-Congo hemorrhagic fever
Rinderpest morbillivirus
Vaccinia virus
Murine norovirus 1
