## Supplementary File 2 for "PanGIA: A Metagenomics Analytical Framework for Routine Biosurveillance and Clinical Pathogen Detection"

### PanGIA Executive Summary Report:

**Sample ID:** PAN-1002-07
**Patient ID:** Unkown patient
**Seq. Run ID:** B
**Seq. Machine ID:** NAMRU
**Analyst:** Hillary Wood
**Sample Matrix:** Other
**Report Reviewed By:**
**Results Validity:**

**Analysis Date:** 09-12-2018 00:00
**Location:** NAMRU-6

**Sample Description:**
NP swab

**The following pathogens were detected with confidence in sample PAN-1002-07:**

| Pathogen | Confidence score | RSNB normalized read count | Confidence Score (Standalone) | Confidence Score (Background) | Coverage | Depth of Coverage | Relative Abundance |
|---|---|---|---|---|---|---|---|
| Escherichia virus MS2 | 0.99 | 21203.12 | 0.99 | NA | 0.96 | 443.76 | 0.9070 |
| Enterobacteria phage phiX174 sensu lato | 0.99 | 2365.18 | 0.99 | NA | 1.00 | 32.85 | 0.0671 |
| Streptococcus pyogenes | 1.00 | 101313.54 | 1.00 | NA | 0.01 | 4.07 | 0.0083 |
| Veillonella parvula | 1.00 | 113738.34 | 1.00 | NA | 0.02 | 3.99 | 0.0082 |
| Streptococcus mitis | 1.00 | 38235.65 | 1.00 | NA | 0.02 | 1.34 | 0.0027 |
| Rothia mucilaginosa | 1.00 | 38673.53 | 1.00 | NA | 0.01 | 1.28 | 0.0026 |
| Streptococcus thermophilus | 1.00 | 13789.67 | 1.00 | NA | 0.02 | 0.54 | 0.0011 |
| Streptococcus salivarius | 0.89 | 11387.24 | 0.89 | NA | 0.03 | 0.39 | 0.0008 |
| Streptococcus parasanguinis | 0.60 | 7536.04 | 0.60 | NA | 0.01 | 0.26 | 0.0005 |
| Streptococcus pneumoniae | 0.63 | 4671.39 | 0.63 | NA | 0.02 | 0.17 | 0.0004 |
| Fusobacterium nucleatum | 0.14 | 3668.42 | 0.14 | NA | 0.01 | 0.12 | 0.0002 |
| Neisseria sicca | 0.19 | 2962.67 | 0.19 | NA | 0.01 | 0.08 | 0.0002 |
| Streptococcus sanguinis | 0.27 | 1954.56 | 0.27 | NA | 0.01 | 0.06 | 0.0001 |
| Neisseria meningitidis | 0.22 | 957.84 | 0.22 | NA | 0.01 | 0.03 | 0.0001 |
| Neisseria lactamica | 0.15 | 893.99 | 0.15 | NA | 0.01 | 0.03 | 0.0001 |
| Streptococcus gordonii | 0.21 | 859.48 | 0.21 | NA | 0.01 | 0.03 | 0.0001 |
| Streptococcus suis | 0.32 | 833.08 | 0.32 | NA | 0.01 | 0.03 | 0.0001 |
| Campylobacter concisus | 0.04 | 682.10 | 0.04 | NA | 0.01 | 0.03 | 0.0001 |
| Streptococcus cristatus | 0.20 | 781.49 | 0.20 | NA | 0.01 | 0.03 | 0.0001 |
| Streptococcus gallolyticus | 0.22 | 893.61 | 0.22 | NA | 0.01 | 0.03 | 0.0001 |
| Neisseria elongata | 0.10 | 772.93 | 0.10 | NA | 0.01 | 0.03 | 0.0001 |
| Streptococcus intermedius | 0.14 | 647.17 | 0.14 | NA | 0.01 | 0.02 | 0.0000 |
| Neisseria weaveri | 0.07 | 609.50 | 0.07 | NA | 0.01 | 0.02 | 0.0000 |
| Streptococcus marmotae | 0.11 | 425.95 | 0.11 | NA | 0.01 | 0.01 | 0.0000 |
| Streptococcus dysgalactiae | 0.07 | 398.66 | 0.07 | NA | 0.01 | 0.01 | 0.0000 |
| Streptococcus anginosus | 0.09 | 382.11 | 0.09 | NA | 0.01 | 0.01 | 0.0000 |
| Streptococcus himalayensis | 0.11 | 380.89 | 0.11 | NA | 0.01 | 0.01 | 0.0000 |
| Streptococcus iniae | 0.06 | 324.72 | 0.06 | NA | 0.01 | 0.01 | 0.0000 |

**Quality Assurance Information:**
Input sequencing reads were trimmed and filtered with the following cut-off settings;
Minimum length: 50 bp
Low complexity ratio: 0.85
Q score > 5
Host Removal: 6155 reads, (0.30%)

| Sample Prepartion | Sequencing | Bioinformatics |
|---|---|---|

| | | | | | | | | |
|---|---|---|---|---|---|---|---|---|
| Sample ID confirmed | NA | | Inter-Run carrayover | | | Exogenous MS2 | Pass | 21234 reads |
| Extraction Quant (DNA) | Fail | 0.0 ng/ul | Index bleedover | NA | % | Exogenous PhiX | Pass | 2370 reads |
| Post-WTA Quant | Pass | 14.3 ng/ul | Analysis Runtime Errors | Fail | No errors at runtime | Mapped Reads | Fail | 0.27% |
| Post-Index PCR Quant | Pass | 5.33 ng/ul | Total Run Yield | Pass | 3.44 GB | Analysis Parameters | | -ams 72 -asl 40 -st standalone -ms 0 -mr 10 -mb 3 -ml 200 -md 0.01 -mrd 0.001 -mc 0.005 -sb -lb /home/edge/edge/thirdParty/pangia/background/QCB_rep1.pangia.json |
| | | | Sequence Run Clusters | NA | M | PanGIA software | | Version 2.4.11.1 BETA |
| | | | Aligned Reads | Fail | 6 % | Database version | | NCBI_genomes_refseq89_Human_GRCh38.p12.fa, NCBI_genomes_refseq89_BAV.fa, NCBI_genomes_refseq89_adds.fa |
| | | | Passed Filter Reads | Pass | 21223859 | | | |

**RESULTS:** --
